# Molecular and Cellular Context Influences SCN8A Variant Function

**DOI:** 10.1101/2023.11.11.566702

**Authors:** Carlos G. Vanoye, Tatiana V. Abramova, Jean-Marc DeKeyser, Nora F. Ghabra, Madeleine J. Oudin, Christopher B. Burge, Ingo Helbig, Christopher H. Thompson, Alfred L. George

## Abstract

Pathogenic variants in *SCN8A*, which encodes the voltage-gated sodium (Na_V_) channel Na_V_1.6, are associated with neurodevelopmental disorders including epileptic encephalopathy. Previous approaches to determine *SCN8A* variant function may be confounded by the use of a neonatal-expressed alternatively spliced isoform of Na_V_1.6 (Na_V_1.6N), and engineered mutations to render the channel tetrodotoxin (TTX) resistant. In this study, we investigated the impact of *SCN8A* alternative splicing on variant function by comparing the functional attributes of 15 variants expressed in two developmentally regulated splice isoforms (Na_V_1.6N, Na_V_1.6A). We employed automated patch clamp recording to enhance throughput, and developed a novel neuronal cell line (ND7/LoNav) with low levels of endogenous Na_V_ current to obviate the need for TTX-resistance mutations. Expression of Na_V_1.6N or Na_V_1.6A in ND7/LoNav cells generated Na_V_ currents that differed significantly in voltage-dependence of activation and inactivation. TTX-resistant versions of both isoforms exhibited significant functional differences compared to the corresponding wild-type (WT) channels. We demonstrated that many of the 15 disease-associated variants studied exhibited isoform-dependent functional effects, and that many of the studied *SCN8A* variants exhibited functional properties that were not easily classified as either gain- or loss-of-function. Our work illustrates the value of considering molecular and cellular context when investigating *SCN8A* variants.

## INTRODUCTION

Pathogenic variants in genes encoding voltage-gated sodium (Na_V_) channels are frequently discovered in individuals with early onset epilepsy, epileptic encephalopathy and related neurodevelopmental disorders (NDD) (1, 2). Determining the functional consequences of Na_V_ channel variants can inform pathophysiological mechanisms and potentially guide precise therapeutic approaches (3, 4). Using the correct molecular context (e.g., species origin, splice isoform) for investigating function of an ion channel variant is vital for an accurate assessment.

Pathogenic variants in *SCN8A*, which encodes Na_V_1.6, have emerged as important causes of neurodevelopmental disorders with typical onset during infancy (5). The earliest discoveries associated epileptic encephalopathy with non-truncating variants having gain-of-function properties (e.g., enhanced persistent current, altered voltage-dependence of activation). Subsequently, *SCN8A* variants were discovered in individuals affected with epilepsy having a wider spectrum of clinical severity as well as NDD without seizures (6).

In mature neurons, Na_V_1.6 is localized to the axon initial segment (AIS) where the channel serves to initiate action potentials (7). The gene undergoes specific alternative splicing events during early development including the in-frame inclusion of one of two distinct versions of exon 5 that encodes a portion of the first voltage-sensing domain (8). Exon 5N dominates during embryonic development and after birth, but around 1 year of age transcripts containing the alternative exon 5A surpass those containing 5N, and the 5A isoform becomes predominant in later childhood through adulthood (9). Importantly, the *SCN8A* reference coding sequence designated as variant 1 (NM_014191) by the National Center for Biotechnology Information (NCBI) includes exon 5N, whereas the sequence including exon 5A is curated as variant 3 (NM_001330260). These annotations guided the early annotation of *SCN8A* variants by genetic testing laboratories such that pathogenic variants in exon 5A were initially overlooked (10).

Previous studies of *SCN8A* variant function used a variety of expression systems including rodent Na_V_1.6 (11-13) or the human channel with exon 5N (neonatal splice isoform), which does not represent the most abundant splice isoform present during most of childhood and older (14-19). In some of these studies, Na_V_1.6 was expressed in a neuronal cell line (ND7/23) that exhibits large endogenous sodium currents. To discriminate between the endogenous and heterologously expressed Na_V_ channels in these cells, many investigators exploited a pore domain mutation to render Na_V_1.6 insensitive to tetrodotoxin (TTX) (11-14, 16-21). However, this strategy could conceivably alter the functional impact of disease-associated variants, especially those located near the TTX interaction site in the channel structure. The heterogeneity in experimental approaches and recombinant Na_V_1.6 channel constructs limits the comparisons of variant dysfunction across laboratories. A uniform experimental approach free from the potential confounds of endogenous currents and second site mutations that also considers the most relevant splice isoform would be valuable.

In this study, we analyzed a series of *SCN8A* variants using automated patch clamp recording of recombinant human Na_V_1.6 heterologously expressed in a derivative of ND7/23 neuronal cells that have low levels of endogenous Na_V_ current. We compared the functional properties of variants in the two isoforms generated by alternative splicing of exon 5 and used the relevant splice isoform to study a subset of variants discovered within exon 5A. Finally, we illustrate the importance of controlling for splice isoform by the functional study of an epilepsy-associated *SCN8A* haplotype with two *de novo* missense variants of uncertain significance within and outside exon 5N. Our findings demonstrate the importance of considering molecular and cellular context when investigating the functional consequences of *SCN8A* variants.

## RESULTS

We initially expressed recombinant WT human Na_V_1.6 in HEK293T cells to evaluate efficiency and fidelity of automated patch clamp as a platform for evaluating the functional consequences of *SCN8A* variants. Results from these initial trials proved unsatisfactory because of inconsistent cell capture and suboptimal membrane seal formation. Therefore, we explored alternative cell lines including the rat dorsal root ganglion neuron-mouse neuroblastoma hybrid ND7/23 cell line (22), which was used successfully by other groups for studying Na_V_1.6 variants (11-13, 16-19, 21). Because ND7/23 cells exhibit a large endogenous fast gating Na_V_ current, previous studies utilized a TTX-resistant mutant form of Na_V_1.6 coupled with recording in the presence of TTX to isolate activity of the transfected channel. We chose an alternative approach designed to eliminate endogenous Na_V_ currents in ND7/23 cells, which obviates the need for TTX and non-native Na_V_1.6 sequences that could confound analysis of disease-associated mutations.

### Generation of a neuronal cell line with minimal Na_V_ current

Previously reported transcriptome analyses (23, 24) and pharmacological studies (25) deduced that Na_V_1.7 was responsible for most of the endogenous Na_V_ current in ND7/23 cells. As additional proof, we transfected ND7/23 cells with siRNA targeting conserved sequences shared by rat and mouse Na_V_1.7 then assessed knockdown success by immunoblotting and manual patch clamp recording. Transient knock-down of Na_V_1.7 eliminated immunodetectable protein and lowered endogenous Na_V_ current substantially (**Fig. 1A,B**). In separate experiments, we failed to detect Na_V_1.6 protein in ND7/23 lysates using an antibody validated against mouse brain (data not shown), indicating that the low level of *Scn8a* mRNA detected in ND/723 cells (23, 24) likely does not contribute to functional Na_V_ channel expression.

**Figure 1.**
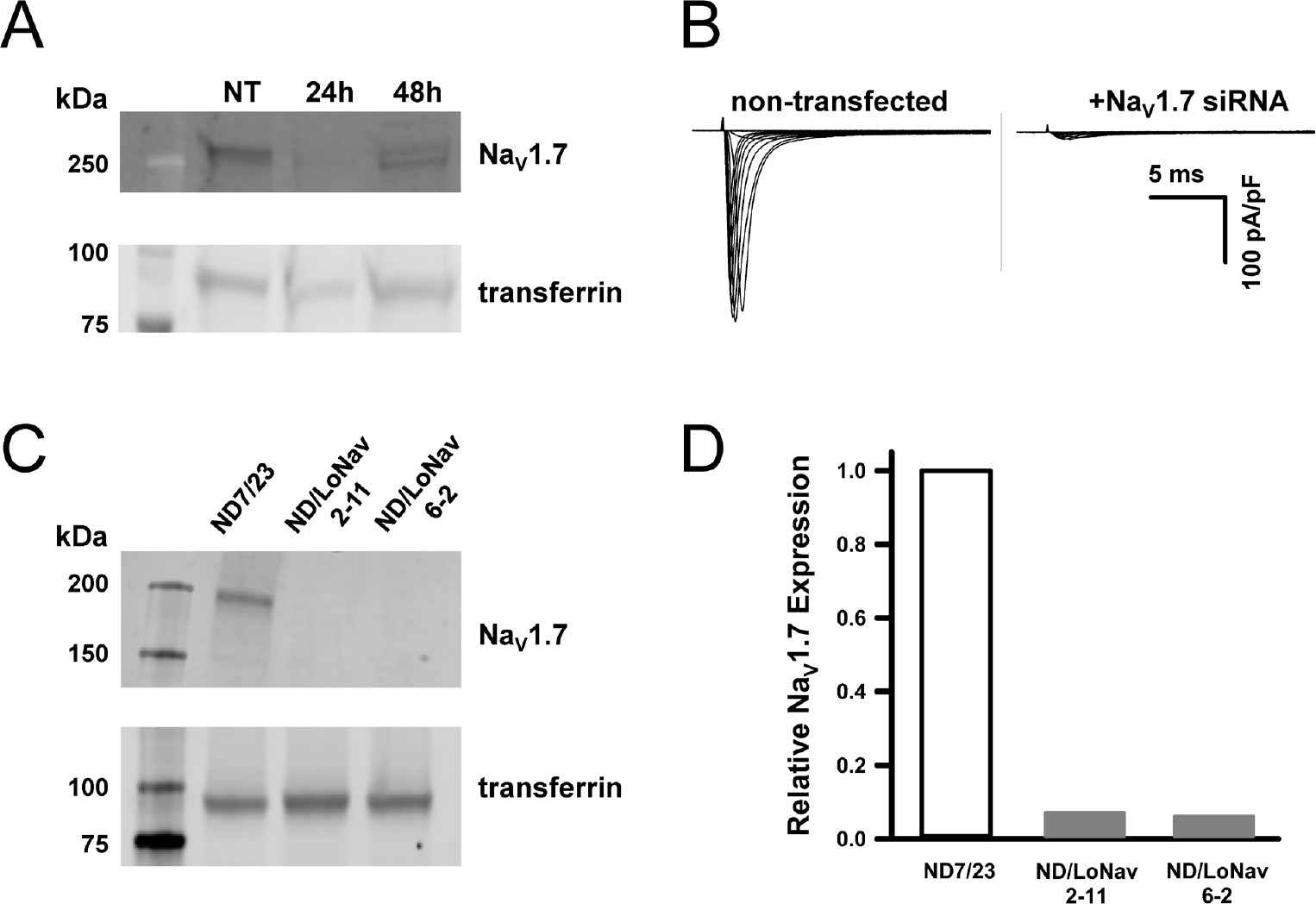
Na_V_1.7 siRNA reduces whole-cell current and Na_V_1.7 protein expression in ND7/23 cells. (A) Immunoblots of Na_V_1.7 and transferrin (loading control) isolated from non-transfected (NT) and Na_V_1.7 siRNA-transfected (24h and 48h post-transfection) ND7/23 cells. (B) Average whole-cell currents from non-transfected (n = 3) and Na_V_1.7 siRNA-transfected (n = 4) ND7/23 cells recorded by manual patch-clamp. (C) Immunoblots of Na_V_1.7 and transferrin isolated from ND7/23 and 2 clonal ND7/LoNav cell lines. (D) Relative Na_V_1.7 protein expression levels from parental ND7/23 cells and 2 clonal ND7/LoNav cell lines.

For stable suppression of endogenous Na_V_ current in ND7/23 cells, we employed CRISPR/Cas9 genome editing to disrupt the coding sequences of rat and mouse Na_V_1.7 (see Methods). An initial round of gene editing yielded clonal lines with multiple frame-shifting deletions in both rat and mouse Na_V_1.7, but a small subpopulation of channel sequences with in-frame deletions were also observed. A second round of gene editing was successful in introducing frame shifts in all detectable Na_V_1.7 transcripts. Two clones with negligible endogenous Na_V_1.7 protein levels (**Fig. 1C,D)** and small endogenous inward currents (peak current density less than 15 pA/pF; **Fig. 2A,B**) were selected. The final clonal cell line (designated ND7/LoNav) was efficiently electroporated with human Na_V_1.6 and proved amenable for high-throughput automated patch clamp analysis.

**Figure 2.**
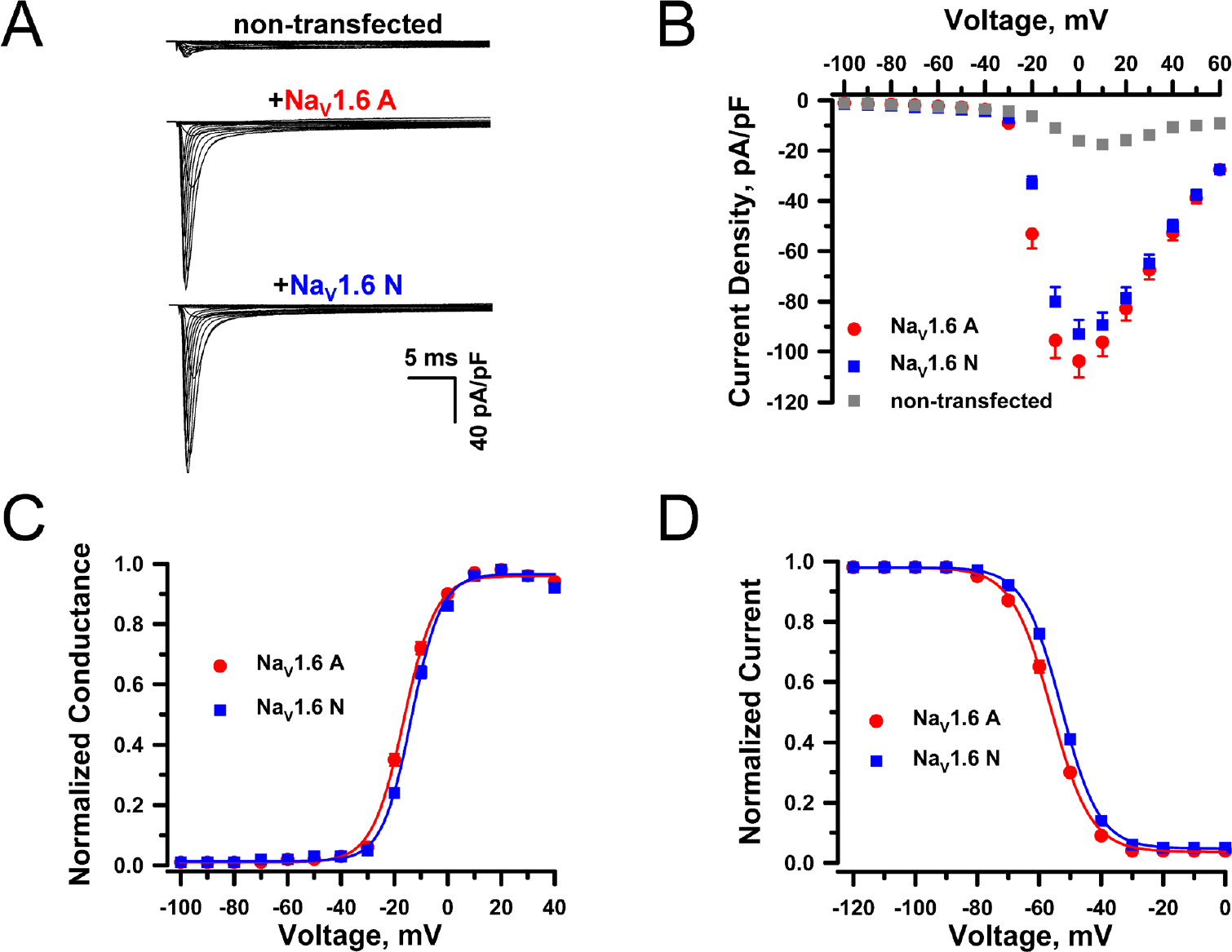
Functional properties of Na_V_1.6N and Na_V_1.6A expressed in ND7/LoNav cells. (A) Average whole-cell currents recorded using automated patch clamp from non-transfected ND7/LoNav cells or transiently transfected with Na_V_1.6A or Na_V_1.6N. Transfection efficiency of WT Na_V_1.6A and Na_V_1.6N averaged 78.3±3.5% and 74.4±5.1%, respectively, and approximately 65% of wells in each 384-well plate exhibited cell capture and formation of stable high resistance (≥0.5 Gohm) membrane seals. (B) Average current density vs voltage measured from non-transfected ND7/LoNav cells (gray squares, n = 103), Na_V_1.6A (red circles, n = 90) or Na_V_1.6N (blue squares, n = 86). (C) Conductance-voltage plots recorded from ND7/LoNav cells expressing Na_V_1.6A (red circles, n = 96) or Na_V_1.6N (blue squares, n = 90). (D) Voltage-dependence of inactivation recorded from ND7/LoNav cells expressing Na_V_1.6A (red circles, n = 139) or Na_V_1.6N (blue squares, n = 142). Quantitative data with statistical comparisons are provided in **Supplemental Dataset S1**.

### Na_V_1.6 splice isoforms exhibit distinct functional properties

We compared the functional properties of adult (Na_V_1.6A) and neonatal (Na_V_1.6N) splice isoforms expressed in ND7/LoNav cells. Cells expressing either Na_V_1.6A or Na_V_1.6N exhibited large, voltage activated inward currents with similar peak current densities, which were ∼6-fold larger and had distinct biophysical properties compared to currents recorded from non-transfected cells (**Fig. 2A,B**; **Supplemental Dataset S1**). Cells expressing either Na_V_1.6 isoform generated whole cell currents with similar inactivation kinetics, but there were significant differences in the voltage dependence of activation and inactivation (**Supplemental Dataset S1**). The Na_V_1.6A isoform exhibited a hyperpolarized and steeper voltage-dependence of activation (**Fig. 2C**) along with a hyperpolarized voltage-dependence of inactivation (**Fig. 2D**) relative to Na_V_1.6N. These results indicate that Na_V_1.6A and Na_V_1.6N exhibit distinct biophysical properties, and suggest that the functional consequences of disease-associated Na_V_1.6 variants may be influenced by the splice isoform.

### Functional consequences of disease-associated *SCN8A* variants

We determined the functional properties of 8 disease-associated *SCN8A* missense variants (Q417P, R850Q, G1475R, R1617L, G1625R, I1631M, N1768D, N1877S) and 2 ultra-rare variants of uncertain significance reported in the ClinVar database (Q713D, G1914S) in both Na_V_1.6A and Na_V_1.6N. Six variants (R850Q, G1475R, R1617L, G1625R, N1768D, N1877S) are recurrent, and 4 (R850Q, G1475R, R1617L, N1768D) were previously investigated to determine mutation-associated functional effects (13, 15-17, 26). Two of the variants were non-recurrent and included Q417P, which was found in a female with intractable infantile spasms and global developmental delay (also reported by (27)), and I1631M discovered in a male with generalized epilepsy with onset at age 5 months followed in later childhood by partial epilepsy and mild intellectual disability. We investigated the functional consequences of three other disease-associated variants discovered in exon 5A (V211R, R223G, I231T) (10) only in Na_V_1.6A. A summary of phenotypes associated with each variant is presented in **Supplemental Table S1**. We also investigated the functional properties of an engineered TTX-resistance mutation (Y371C) (18) in both Na_V_1.6A and Na_V_1.6N. The location of all studied variants on the Na_V_1.6 protein are shown in **Supplemental Fig. S1**. For this study, we analyzed electrophysiological data from 3,265 cells.

Variant-specific effects on channel properties were determined by quantifying differences relative to isoform-matched WT channels that were expressed and recorded in parallel. We analyzed current density, time constant (τ) of fast inactivation at 0 mV, voltage-dependence of activation and inactivation, window current, frequency-dependent loss of channel availability (i.e., current run-down measured at 20 Hz), recovery from fast inactivation, persistent current, and net charge movement during slow voltage ramps. Averaged whole-cell currents normalized to peak isoform-matched WT current are presented in **Supplemental Fig. S2 and Fig. S3** for variants expressed in Na_V_1.6N and Na_V_1.6A, respectively.

Cells expressing most variants exhibited robust current with the exception of R1617L expressed in the Na_V_1.6N splice isoform (**Supplemental Datasets S2 and S3**). Measurable current density was significantly smaller than isoform-matched WT for three variants in the adult isoform and seven in the neonatal isoform. Only one variant (G1475R) exhibited larger current density (**Fig. 3**), and the increase was similar in both isoforms. Activation voltage-dependence varied considerably among this cohort of variants with some exhibiting significantly depolarized or significantly hyperpolarized conductance-voltage relationships (**Fig. 4**). Similarly, the voltage-dependence of inactivation was either significantly depolarized, significantly hyperpolarized or WT-like (**Fig. 5**). The overlap between the activation and inactivation curves illustrated in **Fig. 4B** and **Fig. 5B**, respectively, can be quantified as window current (**Supplemental Fig. S4**), which defines a voltage range in which channels are activated but not inactivated. Most variants exhibited a significantly larger window current consistent with gain-of-function, whereas only one variant (I231T) had a significantly smaller window current.

**Figure 3.**
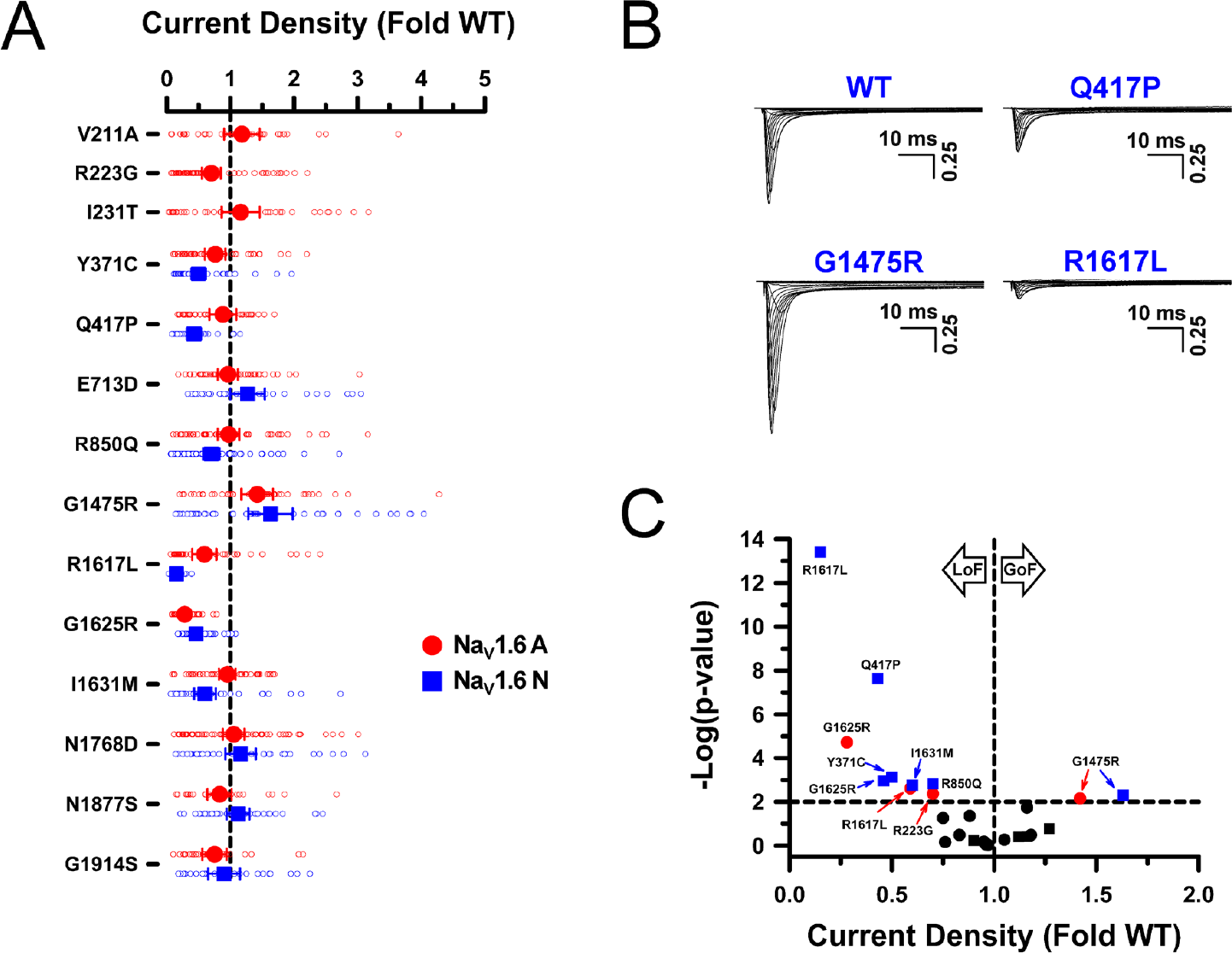
Current density of Na_V_1.6 variants. (A) Peak whole-cell current density for Na_V_1.6 variants and displayed as fold-difference from WT channels recorded in parallel. All individual data points are plotted as open symbols and mean values are shown as larger filled symbols (n = 35-108 per variant). Error bars represent 95% CI. Data from Na_V_1.6A or Na_V_1.6N are indicated as red or blue symbols, respectively. Values to the right or left of the vertical dashed line (normalized WT value) represent current density larger or smaller than WT, respectively. (B) Averaged current traces, normalized to WT peak current density, for WT Na_V_1.6N and select variants with either larger (G1475R) or smaller (Q417P) current density. (C) Volcano plot of mean values highlighting variants with peak current density significantly (P<0.01, horizontal dashed line) different from WT. Symbols to the left of the vertical dashed line denote smaller current (loss-of-function), while symbols to the right indicate larger current (gain-of-function). Black symbols represent variants with no significant difference from WT. Quantitative data with statistical comparisons are provided in **Supplemental Dataset S2** (Na_V_1.6N) and **Supplemental Dataset S3** (Na_V_1.6A).

**Figure 4.**
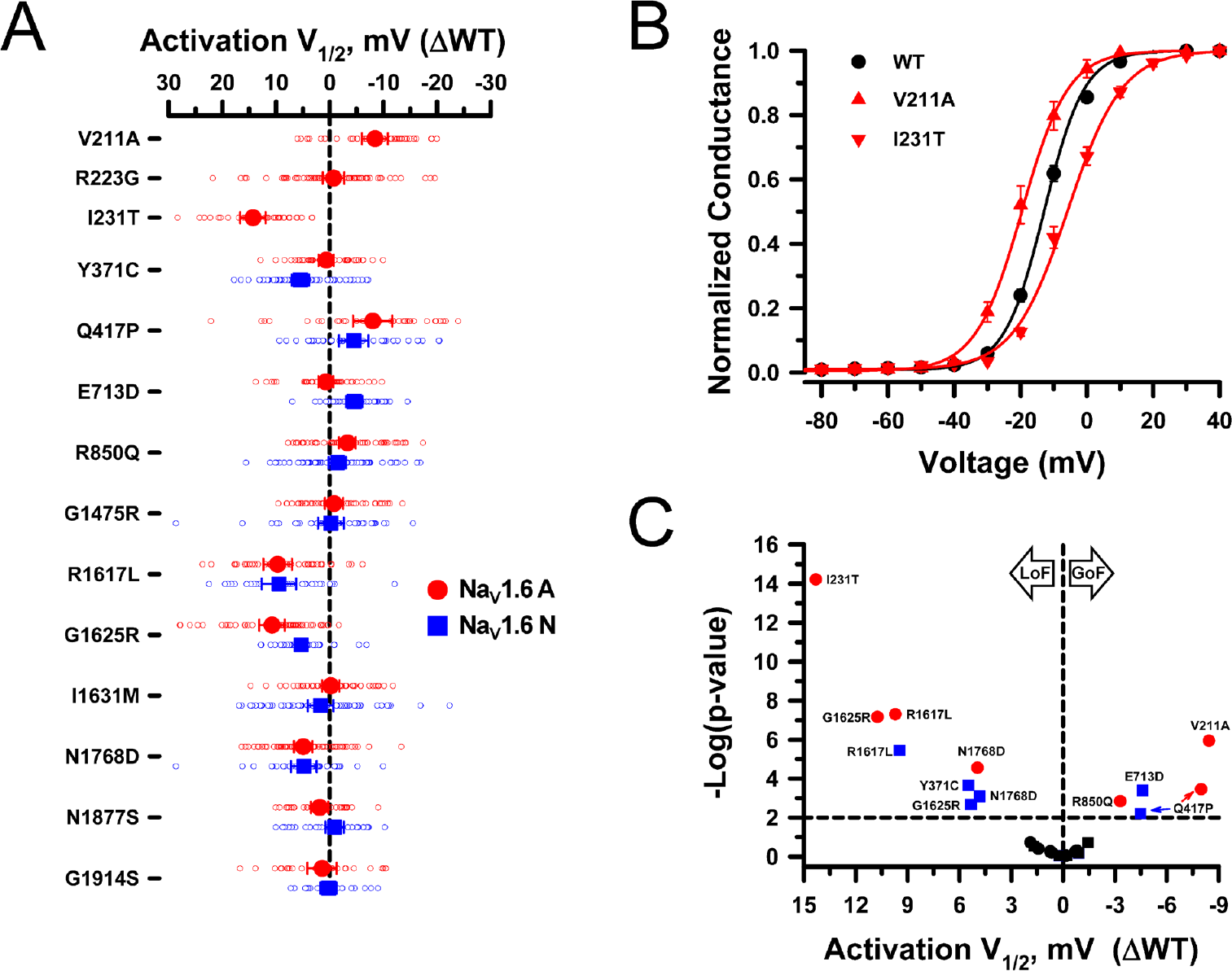
Voltage-dependence of activation for Na_V_1.6 variants. (A) Averaged voltage-dependence of activation V½ obtained from fitting the data for each variant-expressing cell and plotted as difference (ΔV½ in mV) from the averaged V½ for WT channels recorded in parallel. All individual data points are plotted as open symbols and mean values are shown as larger filled symbols (n = 22-77 per variant). Error bars represent 95% CI. Data from Na_V_1.6A or Na_V_1.6N are indicated as red or blue symbols, respectively. Values to the right or left of the vertical dashed line (no difference from WT) indicate hyperpolarized (gain-of-function) or depolarized (loss-of-function) activation V½, respectively. (B) Conductance-voltage relationships for select variants expressed in Na_V_1.6A (red lines) illustrating hyperpolarized (V211A) or depolarized (I231T) shifts in activation V½ relative to WT channels (black line) recorded in parallel. (C) Volcano plot of mean values highlighting variants with significantly different (P<0.01, horizontal dashed line) activation V½. Symbols to the left of the vertical dashed line denote depolarized V½ values (loss-of-function), while symbols to the right indicate hyperpolarized V½ values (gain-of-function). Black symbols represent variants with no significant difference from WT. Quantitative data with statistical comparisons are provided in **Supplemental Dataset S2** (Na_V_1.6N) and **Supplemental Dataset S3** (Na_V_1.6A).

**Figure 5.**
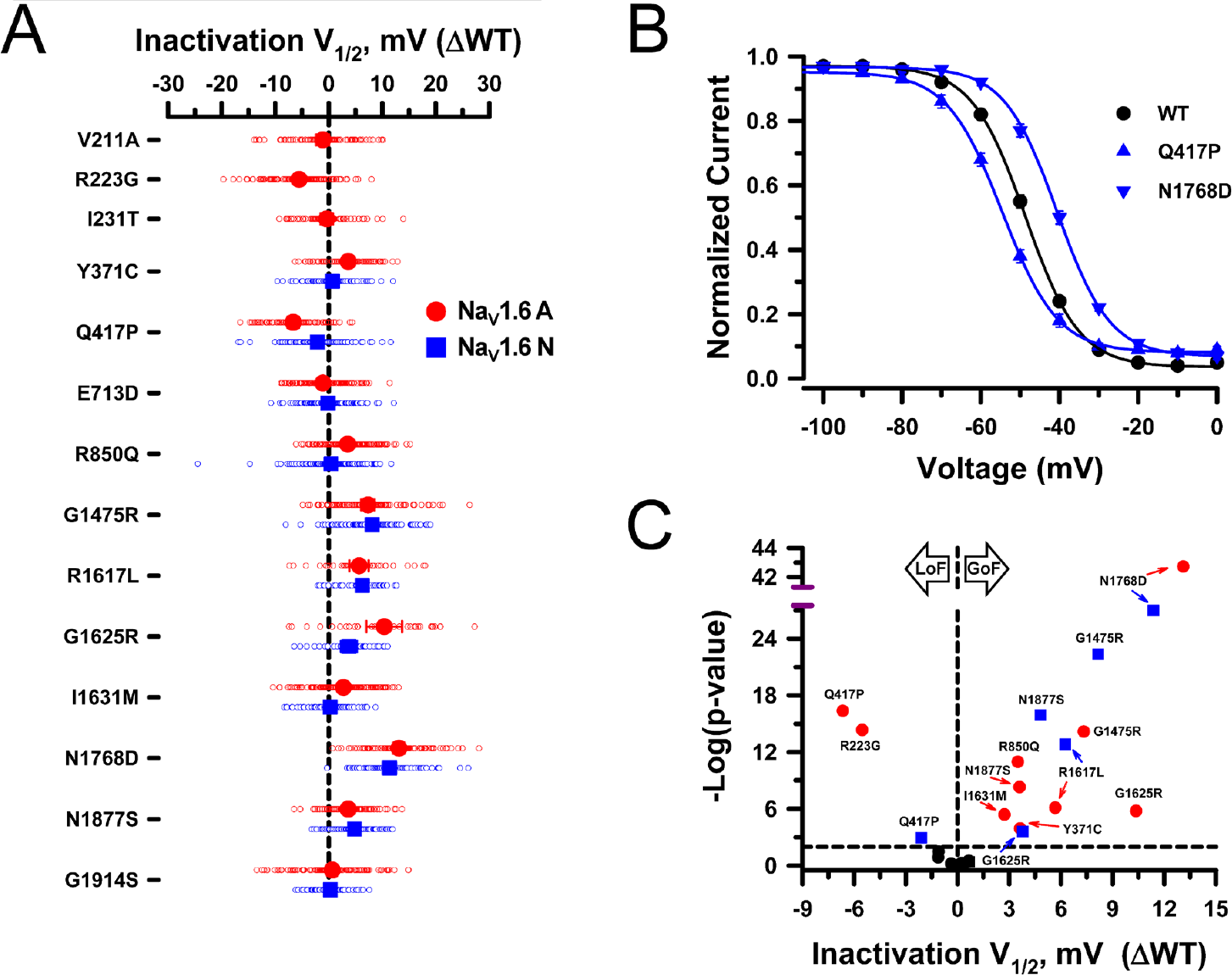
Voltage-dependence of inactivation for Na_V_1.6 variants. (A) Averaged voltage-dependence of inactivation V½ obtained from fitting the data for each variant-expressing cell and plotted as difference (ΔV½ in mV) from the averaged V½ for WT channels recorded in parallel. All individual data points are plotted as open symbols and mean values are shown as larger filled symbols (n = 29-142 per variant). Error bars represent 95% CI. Data from Na_V_1.6A or Na_V_1.6N are indicated as red or blue symbols, respectively. Values to the right or left of the vertical dashed line (no difference from WT) indicate depolarized (gain-of-function) or hyperpolarized (loss-of-function) inactivation V½, respectively. (B) Steady-state inactivation curves for select variants expressed in Na_V_1.6N (blue lines) illustrating hyperpolarized (Q417P) or depolarized (N1768D) inactivation V½ relative to wild type channels (black line) recorded in parallel. (C) Volcano plot of mean values highlighting variants with significantly different (P<0.01, horizontal dashed line) inactivation V½. Symbols to the left of the vertical dashed line denote hyperpolarized V½ values (loss-of-function), while symbols to the right indicate depolarized V½ values (gain-of-function). Black symbols represent variants with no significant difference from WT. Quantitative data with statistical comparisons are provided in **Supplemental Dataset S2** (Na_V_1.6N) and **Supplemental Dataset S3** (Na_V_1.6A).

Inactivation kinetics (measured at 0 mV), persistent current and ramp current amplitudes were also variable with notable slowing of inactivation time course and greater persistent or ramp current for multiple variants (**Fig. 6, 7**), which are consistent with gain-of-function. Only R223G exhibited smaller persistent or ramp current then WT. Persistent current was most prominent for the recurrent epilepsy-associated variant N1768D in agreement with previous reports (11, 14). The kinetics observed for recovery from inactivation (**Supplemental Fig. S5**) and frequency-dependent current rundown (**Supplemental Fig. S6**) illustrate differences for some variants that are consistent with loss-of-function. The variants with the slowest recovery from inactivation were N1768D and Q417P expressed in either Na_V_1.6N or Na_V_1.6A, and this correlated with variable degrees (5-13%) of frequency-dependent current rundown measured at 20 Hz.

**Figure 6.**
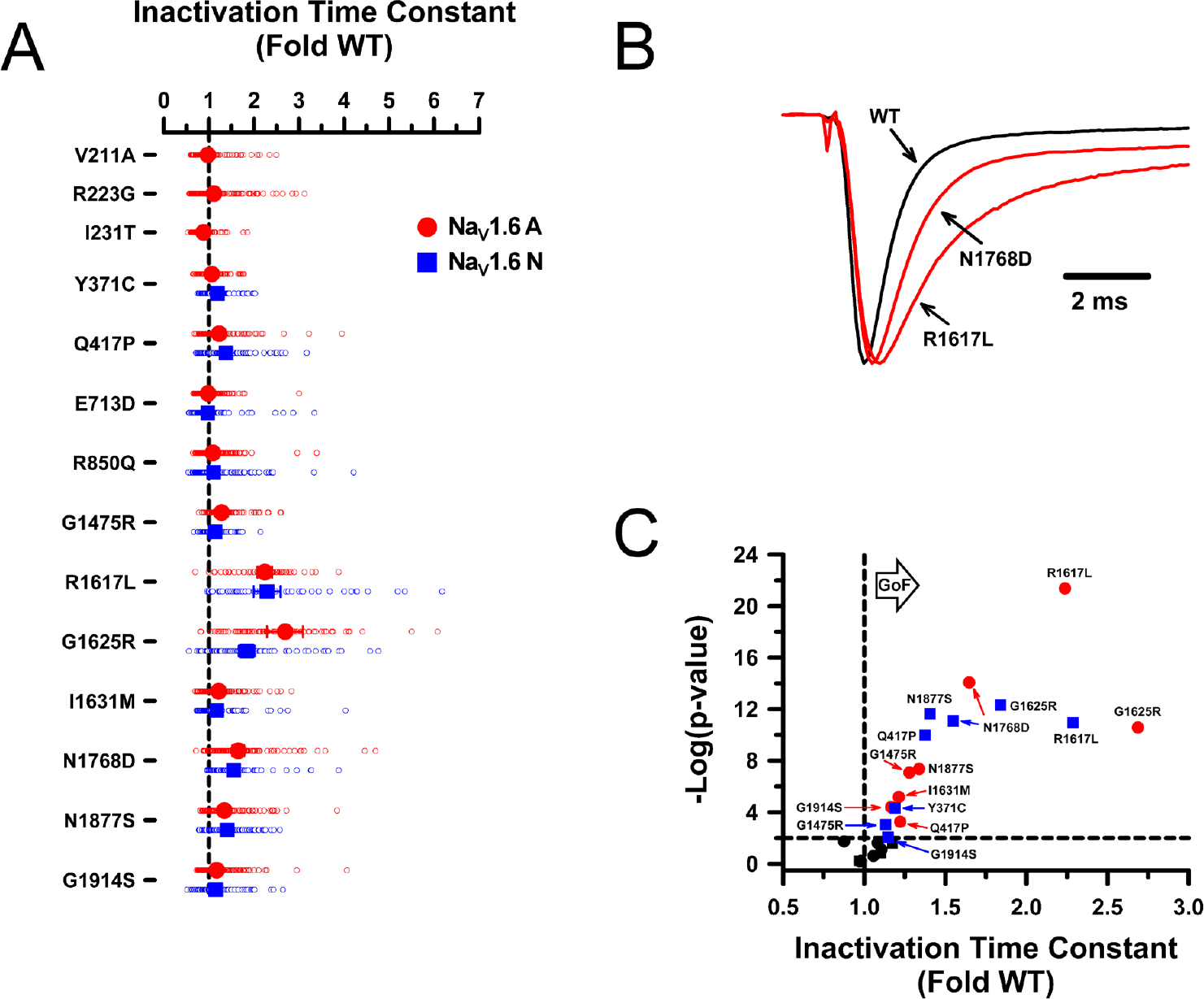
Inactivation kinetics for Na_V_1.6 variants. (A) Averaged inactivation time constants measured at 0 mV obtained by fitting current decay for each variant cell with a single exponential function and expressed as a ratio to the averaged WT channel value recorded in parallel. All individual data points are plotted as open symbols and mean values are shown as larger filled symbols (n = 57-160 per variant). Error bars represent 95% CI. Data from Na_V_1.6A or Na_V_1.6N are indicated as red or blue symbols, respectively. Values to the right or left of the vertical dashed line (average normalized WT value) indicate slower (gain-of-function) or faster (loss of function) inactivation kinetics, respectively. (B) Averaged traces recorded at 0 mV, normalized to the peak current density for select variants illustrating faster (V211A) or slower (N1768D) inactivation. (C) Volcano plot of mean values highlighting variants with significantly different (P<0.01, horizontal dashed line) inactivation time constants. Symbols to the right of the vertical dashed line represent slower inactivation kinetics (gain-of-function). No variants exhibited significantly faster inactivation. Black symbols represent variants with no significant difference from WT. Quantitative data with statistical comparisons are provided in **Supplemental Dataset S2** (Na_V_1.6N) and **Supplemental Dataset S3** (Na_V_1.6A).

**Figure 7.**
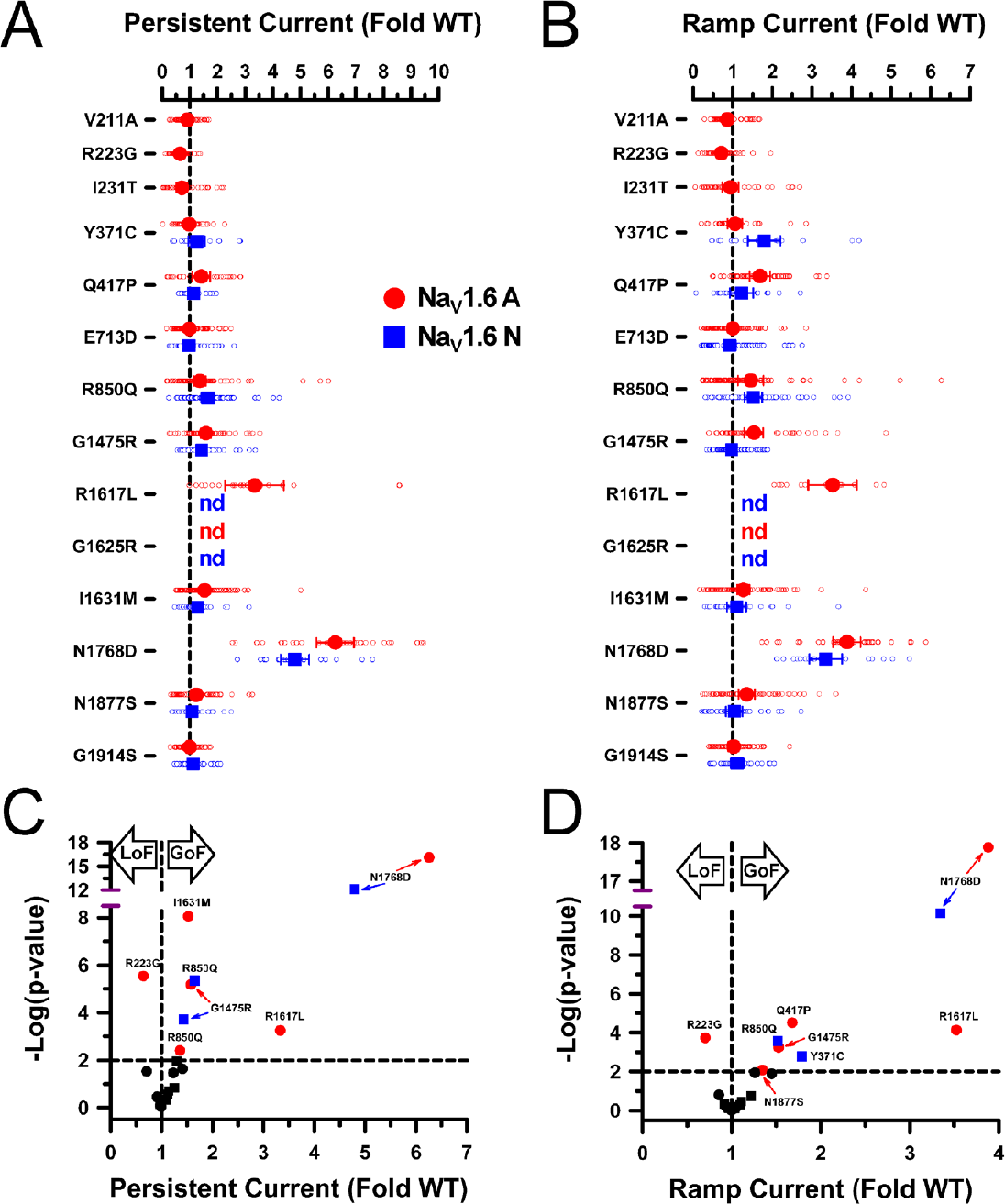
Persistent and ramp currents for Na_V_1.6. (A,B) Average persistent current amplitude (A) or net charge movement during a depolarizing voltage ramp (B) for variants displayed as fold-differences from WT channels recorded in parallel. All individual data points are plotted as open symbols and mean values are shown as larger filled symbols (n = 9-94 per variant). Values for some variants were not determined (nd) because whole-cell current density was too small. Error bars represent 95% CI. Data from Na_V_1.6A or Na_V_1.6N are indicated as red or blue symbols, respectively. Values to the right or left of the vertical dashed lines (average normalized WT values) indicate larger (gain-of-function) or smaller (loss of function) currents, respectively. (C,D) Volcano plots of mean values highlighting variants with significantly different (P<0.01, horizontal dotted line) levels of persistent current (C) or ramp current (D) compared to WT channels. Symbols to the left of the vertical dashed lines denote smaller current levels (loss-of-function), while symbols to the right indicate larger current levels (gain-of-function). Black symbols represent variants with no significant difference from WT. Quantitative data with statistical comparisons are provided in **Supplemental Dataset S2** (Na_V_1.6N) and **Supplemental Dataset S3** (Na_V_1.6A).

Our investigation of the functional consequences of *SCN8A* using the two alternatively spliced versions of the channel revealed several variants with significant isoform-dependent properties. For a given variant, the magnitude of difference compared with the isoform-matched WT channel can be greater in either Na_V_1.6N or Na_V_1.6A (summarized in radar plots, **Supplemental Fig. S7**). For example, the recurrent variant G1625R exhibits a greater degree of inactivation slowing and larger shifts in activation and inactivation V_½_ in Na_V_1.6A, whereas the degree of dysfunctional was less severe in the neonatal isoform.

Our results demonstrate that disease-associated Na_V_1.6 variants exhibited various biophysical defects consistent with either gain-of-function (e.g., slower and less complete inactivation, hyperpolarized voltage-dependence of inactivation) or loss-of-function (depolarized voltage-dependence of activation, slower recovery from inactivation). However, opposing functional properties are found for specific variants and this confounds the assignment of an overall effect using a simple binary classification scheme. To provide a visual summary of the functional properties assessed for each variant expressed in either Na_V_1.6N or Na_V_1.6A, we generated heat maps that scaled each measured parameter along a functional axis from loss to gain (**Supplemental Fig. S8**). The heat maps illustrate that most of the Na_V_1.6 variants exhibit complex patterns of dysfunction, a pattern also observed for *SCN2A* (Na_V_1.2) variants (28). The two *SCN8A* VUS studied exhibited minimal to no functional perturbations.

### Functional effects of a TTX-resistant variant

An advantage of using ND7/LoNav cells is the ability to express a Na_V_ channel in a neuronal cell environment without requiring a pharmacological strategy to negate endogenous current. Because engineering TTX-resistance has been widely used to distinguish Na_V_1.6 current from endogenous Na_V_ current in the original ND7/23 cell line, we examined functional properties of the Y371C TTX-resistant variant (16, 18, 19) expressed in ND7/LoNav cells. Expression of Y371C gave rise to measurable voltage-dependent Na_V_ currents in both Na_V_1.6N and Na_V_1.6A, but the current density in the neonatal isoform was significantly smaller than cells expressing the isoform-matched WT channel (**Fig. 3A**). In addition, the voltage-dependence of activation was significantly shifted for Na_V_1.6N-Y371C (**Fig. 4A**), and recovery from inactivation was significantly slower (**Supplemental Fig. S5**). The adult isoform was less affected by Y371C, but there were significant differences in the voltage-dependence of inactivation (**Fig. 5A**) and recovery from fast inactivation (**Supplemental Fig. S5**). Our findings indicate that Y371C impacts function of Na_V_1.6 in our experimental system.

### Value of studying variants in Na_V_1.6 splice isoforms

To emphasize further the importance of studying disease-associated variants the most relevant Na_V_1.6 splice isoform, we demonstrated the functional consequences of a complex *SCN8A* genotype discovered in a female infant with early-onset epileptic encephalopathy. The affected individual exhibited focal seizures and infantile spasms beginning at age 3 months, which were largely refractory to multiple drug treatments. She later developed tonic seizures, hypotonia and cortical visual impairment. Clinical genetic testing identified two *de novo* missense variants of uncertain significance (c.431C>G, p.T144S; c.649T>C, p.S217P). Subsequent long-read genomic sequence analysis identified a third synonymous variant (c.660C>G, p.R220R) and demonstrated that all three variants were present on the same *SCN8A* allele. Importantly, both S217P and R220R are located within exon 5N, whereas T144S was in a neighboring exon not subject to alternative splicing. Because of the presumed developmentally-regulated exclusion of exon 5N and uncertainty about which variant was pathogenic, we investigated the functional properties of the compound T144S/S217P/R220R genotype in neonatal Na_V_1.6N, and the single T144S variant in the Na_V_1.6A isoform (**Fig. 8**).

**Figure 8.**
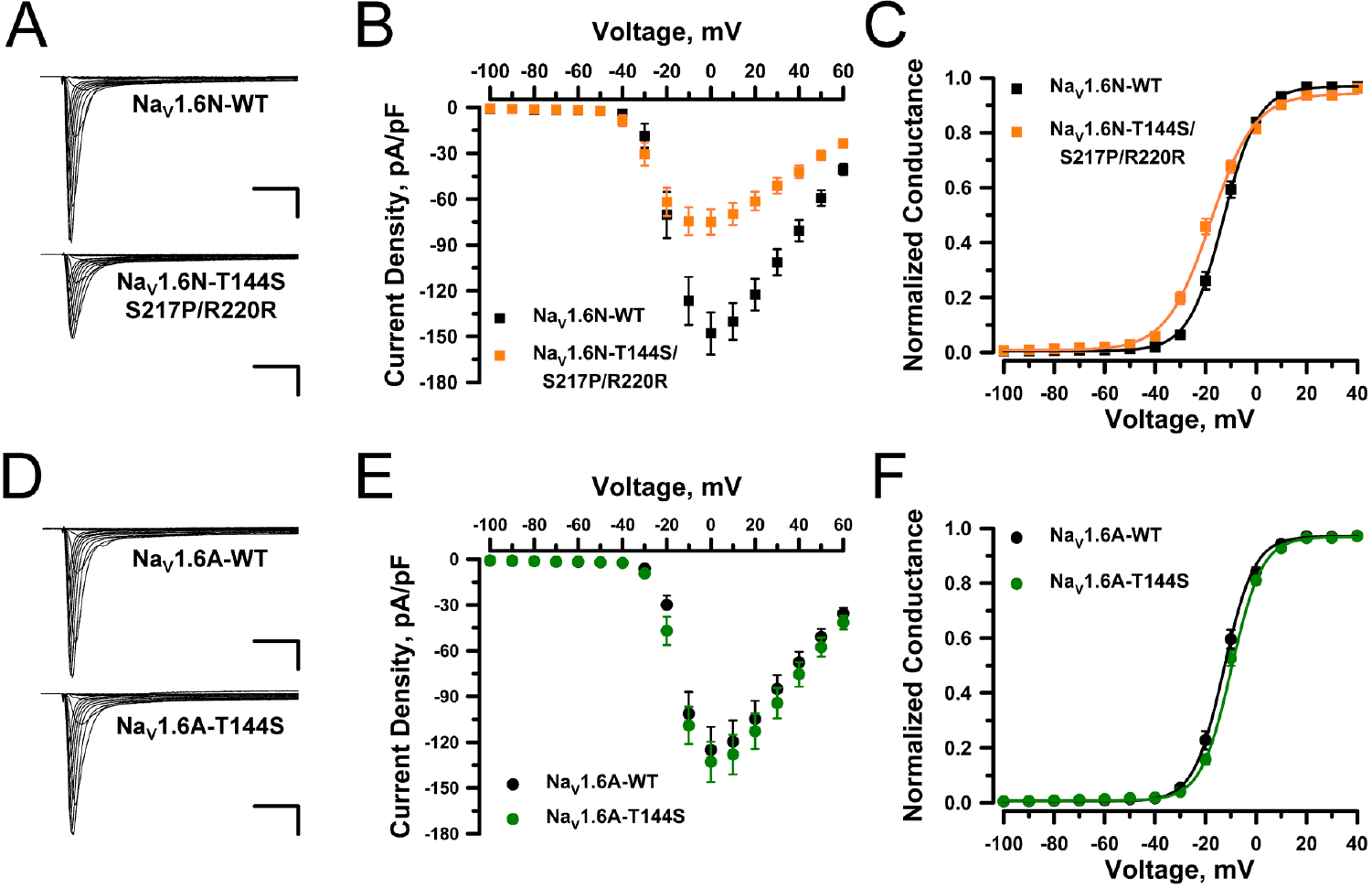
Functional properties of a complex *SCN8A* genotype of uncertain significance. (A) Averaged whole-cell current density recorded from ND7/LoNav cells expressing WT or variant Na_V_1.6N channels (n = 71-78 per variant). Scale bars represent 5 ms (horizontal) and 25 pA/pF (vertical). (B) Current-voltage plots comparing WT (black symbols and lines) and variant (orange symbols and lines) Na_V_1.6N channels. Current amplitudes were normalized to cell capacitance to determine current density. (C) Conductance-voltage plots for WT (black symbols and lines) and variant (orange symbols and lines) Na_V_1.6N channels. (D) Averaged whole-cell current density recorded from ND7/LoNav cells expressing WT or variant Na_V_1.6A channels (n = 51-75 per variant). Scale bars represent 5 ms (horizontal) and 25 pA/pF (vertical). (E) Current-voltage plots comparing WT (black symbols and lines) and variant (green symbols and lines) Na_V_1.6A channels. Current amplitudes were normalized to cell capacitance to determine current density. (F) Conductance-voltage plots for WT (black symbols and lines) and variant (green symbols and lines) Na_V_1.6A channels. Quantitative data with statistical comparisons are provided in **Supplemental Dataset S2** (Na_V_1.6N) and **Supplemental Dataset S3** (Na_V_1.6A). Error bars in panels B,C,E, and F represent 95% CI.

Functional analysis of the triple variant in Na_V_1.6N revealed that compared with the WT channel the compound variant exhibited significantly smaller whole-cell currents, significantly hyperpolarized shifts in the voltage-dependence of activation and inactivation, slower recovery from inactivation, and larger persistent and ramp currents (**Fig. 8A-C, Supplemental Dataset S2**). By contrast, T144S expressed in the Na_V_1.6A isoform exhibited mostly WT-like properties with the exception of slightly slower inactivation kinetics and modestly larger ramp current and window current (**Fig. 8D-E, Supplemental Dataset S3**). Although only the triple variant channel had altered voltage-dependence of inactivation, both variants exhibited slower time course of inactivation measured at 0 mV, which was greater for the compound variant expressed in Na_V_1.6N. We concluded that the compound variant exhibits a predominantly gain-of-function phenotype in the neonatal splice isoform and is a likely pathogenic driver of the clinical phenotype. This also suggests that suppression of exon 5N incorporation into mature *SCN8A* mRNA transcripts may be a therapeutic strategy in this case.

## DISCUSSION

In this study, we demonstrated the importance of considering alternative splicing when assessing disease-associated *SCN8A* variants, and validated two new experimental strategies for accomplishing this task (creation of a neuronal cell line without endogenous sodium currents and use of automated patch clamp recording). In combination, these approaches provide the means for greater physiological relevance and higher throughput that will help standardize efforts to determine the functional consequences of *SCN8A* variants, which is important for assessing variant pathogenicity, ascertaining molecular pathogenesis, and for screening therapeutic agents (21). In particular, automated patch clamp recording is increasingly used to assess the functional consequences of human ion channel variants at a scale difficult to achieve with traditional electrophysiological methods (28-38).

A focus of our study was on comparing the functional properties of WT and variant Na_V_1.6 expressed in two distinct splice isoforms of the channel (Na_V_1.6N, Na_V_1.6A). Human brain Na_V_ channel genes undergo developmentally regulated alternative splicing including a well-characterized event involving two distinct versions of the fifth coding exon that encode portions of the first voltage-sensing domain (8). Transcriptome profiling of human brain indicates that Na_V_1.6N is the most abundant *SCN8A* transcript before birth whereas Na_V_1.6A becomes the major splice isoform by around age 1 year (9) with a transition sometime during infancy.

At the typical age of onset of neurological symptoms in *SCN8A*-related disorders (4-6 months), there is a mix of splice isoforms containing either exon 5A or 5N (9). Published studies on the functional consequences of *SCN8A* variants only use cDNA constructs representing rodent Na_V_1.6 (splice isoform unclear) (11-13) or the neonatal splice isoform of the human channel (14-19). It is conceivable that pathogenic *SCN8A* variants disrupt the timing of alternative splicing in one direction or the other. Our study considered both splice isoforms to ensure we captured the most physiologically relevant molecular context, and determined that the functional properties for some variants are isoform dependent.

*SCN8A* exons 5N and 5A encode proteins that differ by two amino acids at positions 207 (5A: isoleucine; 5A: valine) and 212 (5N: asparagine; 5A: aspartate), and these differences are sufficient to affect the functional properties of the respective channel splice isoforms (**Fig. 2; Supplemental Dataset S1**. The neurophysiological consequences of this alternatively splicing event for *SCN8A in vivo* are unknown, but the analogous splicing event in mouse Na_V_1.2 correlates with greater cortical pyramidal neuron excitability beginning after postnatal day 3 that persists into adulthood (39) and a computational model of a human cortical pyramidal neuron exhibits higher firing frequency when Na_V_1.2A is incorporated (40). Based on our electrophysiological observations, Na_V_1.6N exhibits depolarized voltage-dependence of both activation and inactivation that have the potential of affecting neuronal excitability. Prior studies suggested that shifts in Na_V_1.6 activation voltage-dependence have a greater impact on neuronal firing than similar shifts in the voltage-dependence of inactivation (16), and we could speculate that Na_V_1.6N expression renders immature neurons less excitable.

Because we considered the molecular context for *SCN8A* variants to be relevant to understanding their functional consequences, we investigated the functional properties of variants located exclusively in either exon 5A or 5N using the relevant splice isoform. One variant (R223G) was previously studied in a TTX-resistant version of human Na_V_1.6N expressed in ND7/23 cells (20), and findings from that study differ from our data on human Na_V_1.6A-R223G expressed in ND7/LoNav cells (20). Specifically, the previous study demonstrated hyperpolarized activation, WT-like inactivation voltage-dependence and faster recovery from inactivation, whereas our data indicate WT-like activation voltage-dependence, hyperpolarized inactivation voltage-dependence and slower recovery from inactivation. Two other variants (R850Q, G1475R) previously studied in TTX-resistant Na_V_1.6N also exhibited functional differences with our study using the human Na_V_1.6A isoform. The previous work on R850Q highlighted large persistent current and hyperpolarized voltage-dependence of inactivation (17), which we also observed, but our findings differed in the impact on activation voltage-dependence. Similarly, prior work on G1475R in human TTX-resistant Na_V_1.6N (16, 18) differed from our work with Na_V_1.6A in current density, persistent current and ramp current. Not all variants, including the recurrent variant N1768D, exhibit isoform-dependent properties.

The value of studying variants in the correct molecular context was also illustrated by our investigation of a complex genotype with three in-phase variants of uncertain significance discovered in an infant with severe epileptic encephalopathy. This work allowed us to demonstrate which genotype was most likely to be pathogenic, and provided data supporting the potential therapeutic value of induced exon 5 splice switching (41).

In addition to considering the relevant splice isoform, we also developed a new cellular platform for investigating *SCN8A* variants that exploits a neuronal cell environment without requiring a pharmacological intervention to isolate human Na_V_1.6 current. We developed ND7-LoNav cells by genetically inactivating endogenous rodent Na_V_1.7 resulting in a neuron-derived cell line without appreciable background sodium current. Use of these cells obviates the need for a second site mutation to render human Na_V_1.6 TTX-resistant (14, 16-19). Our novel approach avoids the potential confounding effects of the TTX-R mutation, which we demonstrated have significant functional differences from WT channels in either Na_V_1.6N or Na_V_1.6A. Other studies have shown that a similar TTX-R mutation in Na_V_1.3 also exhibits dysfunctional properties (42). We raise concern that some prior findings may be confounded by extra pore domain mutations.

One goal of determining the functional consequences of *SCN8A* variants is to aid in building genotype-phenotype correlations. Such correlations are valuable for understanding differences in pathophysiological mechanisms and may help guide pharmacological therapy such as the use of Na_V_ channel blocking anti-seizure medications, which would be most useful in the setting of gain-of-function variants. The majority of variants we studied exhibited multiple functional disturbances that individually are consistent with either gain- or loss-of-function, but the net effect of these effects can be challenging to determine without additional experimental work. For some variants such as N1768D, enhanced persistent current combined with depolarized voltage-dependence of inactivation are likely drivers of elevated seizure susceptibility. However, even this widely accepted gain-of-function variant exhibits other functional properties that are consistent with loss-of-function including depolarized voltage-dependence of activation (**Fig. 4**), slower recovery from inactivation (**Supplemental Fig. S5**) and a greater tendency for frequency-dependent loss of activity (**Supplemental Fig. S6**). Complementary experimental approaches with computational simulation of neuronal action potentials (43) or dynamic action potential clamp (44, 45) may be valuable next steps.

In summary, we standardized a new approach for evaluating the functional consequences of *SCN8A* variants that exploits the higher throughput achievable with automated patch clamp recording and obviates the need for second site TTX-resistance mutations through use of a novel neuronal cell line with low levels of endogenous Na_V_ current. Our findings demonstrate that developmentally regulated alternative splicing of exon 5 influences variant function and emphasize the importance of studying variants in physiologically relevant splice isoform.

## METHODS

### Study approval

For unpublished *SCN8A* variants, we obtained informed consent from parents to allow publication of brief de-identified clinical descriptions and genetic information using a method approved by the Children’s Hospital of Philadelphia Institutional Review Board.

### Long-read SCN8A genomic sequencing

Cheek swab genomic DNA was obtained from a proband carrying two *de novo SCN8A* missense variants and from both parents. To determine the phase of the two variants in the proband, long-range genomic PCR was performed under dilute conditions with long extension times and Phusion High Fidelity Polymerase (ThermoFisher) using primers targeting an amplicon of ∼2.6 kb encompassing both variants (forward: CTCTTCTGTGCTTCACCTTTCTCTAGC; reverse: CCTATCCCAACACCTAACACCAACC). Samples were analyzed for quantity and quality using UV-Vis spectrometry and FemtoPulse pulsed-field capillary electrophoresis, followed by standard SMRTbell PacBio library preparation. Library quality control was done using Qubit fluorometric analysis.

Long-read sequencing of the 3 libraries was performed using a single flow cell on a PacBio Sequel v3r9 (Pacific Biosciences, Menlo Park, CA) at the Massachusetts Institute of Technology Biomicro Center using PacBio Circular Consensus Sequencing to improve base-calling accuracy. Sequence reads from the proband sample were first filtered for lengths in the 2500-2700 bp range. Sequences were then filtered for the presence of four 21-bp “anchor” segments (A, B, C, D) arranged AV_1_B to CV_2_D. Anchor segments were defined as 21 bp reference genomic sequences located immediately upstream or downstream of each variant (V_1_, V_2_), with no filter on the identity of the base at the V_1_ or V_2_ positions. After discovery of a third synonymous variant (V_3_) described below, the filter for the D anchor was adjusted to allow any base at the location of V_3_ in this anchor. This filtering yielded 73,522 sequences. Sequences were scored for the presence of the variant (V) or reference (R) allele at the V_1_ or V_2_ positions, yielding 46% V/V, 40% R/R, 7% V/R, and 7% R/V sequences, indicating a relatively low representation of chimeric products, and indicating that the two variants are in *cis*. Similar analyses of the parental samples yielded ∼99% reference allele at both variant positions, confirming that both variants arose *de novo* in the proband. Visual inspection of multiple sequence alignments of proband amplicon sequences also revealed the presence of a third synonymous variant, V_3_: g.52082587C>G (chr12, hg19) located 11 bp downstream of V_2_. This variant was observed in >99% of proband sequences containing V_2_ and in virtually none of the sequences containing the reference allele at this position, confirming that it is in *cis* relative to V_2_ (and V_1_ by inference). The V_3_ variant was not observed in the parental sequences, confirming that it is also arose *de novo* in the proband.

### Plasmids and mutagenesis

Plasmids encoding human Na_V_1.6 splice isoforms annotated by NCBI as variant 1 (NM_014191; neonatal-expressed, Na_V_1.6N) and variant 3 (NM_001177984; adult-expressed, Na_V_1.6A) were rendered stable in bacteria by inserting small introns at the exon 14-15 and 22-23 junctions as described previously (22). Plasmids included an IRES2 element followed by the reading frame for the red fluorescent protein mScarlet to enable determination of transfection efficiency. Both plasmids are available from AddGene (Na_V_1.6N, 162280; Na_V_1.6A 209411). *SCN8A* variants were introduced into wild-type (WT) Na_V_1.6A or Na_V_1.6N using PCR mutagenesis with Q5^®^ Hot Start High-Fidelity 2X Master Mix (New England Biolabs) as described previously (22). Primers were designed for each mutation using custom software to have a minimum 5’ overlap of 20 bp and a predicted melting temperature (Tm) of 60°C (**Supplemental Table S2**). All recombinant plasmids were sequenced in their entirety using nanopore-based sequencing (Primordium Labs, Arcadia, CA, USA) to confirm the presence of the desired modifications and the absence of inadvertent mutations.

### Cell culture

ND7/23 (Sigma-Aldrich, St. Louis, MO, USA) and ND7/LoNav (see below) cells were grown at 37°C with 5% CO_2_ in Dulbecco’s modified Eagle’s medium (DMEM) supplemented with 10% fetal bovine serum (ATLANTA Biologicals, Norcross, GA, USA), 2 mM L-glutamine, and penicillin (50 units/ml)-streptomycin (50 µg/ml). Unless otherwise stated, all tissue culture media was obtained from ThermoFisher Scientific (Waltham, MA, USA).

### Generation of ND7/LoNav cells

Transient knockdown of Na_V_1.7 mRNA was done using short interfering RNA (siRNA) targeting both mouse and rat channel transcripts (Cat# 4390771, ID: s134909; Thermo Fisher Scientific Ambion, Waltham, MA, USA). Permanent knockout of endogenous Na_V_1.7 in ND7/23 cells was achieved using CRISPR/Cas9 genome editing with plasmid pD1301-AD (pCMV-Cas9-2A-GFP) encoding a guide RNA (gRNA) sequence GTTACTGCTGCGCCGCTCCC targeting both mouse *Scn9a* (chr2:66540435-66540454) and rat *Scn9a* (chr3:59260717-59260736). Genome editing vectors were synthesized by ATUM (Newark, CA, USA). ND7/23 cells were transfected with FuGENE® 6 Transfection Reagent (Promega, Madison, WI, USA) according to the manufacturer’s instructions. Two days after transfection, cells were flow-sorted and green fluorescent cells with the top 50% intensity were isolated for single-clone selection. Sequencing of PCR amplicons encompassing the edited regions that were subcloned into the TOPO-TA vector (Invitrogen) revealed a subset with in-frame deletions in mouse *Scn9a*. An additional round of CRISPR/Cas9 genome editing was performed on one clonal cell line to further disrupt the coding region of endogenous *Scn9a* using recombinant pX458 plasmid (pSpCas9-2A-GFP; Addgene #48138) and a gRNA targeting the in-frame deletion (*Scn9a*, CTATTTGTACCCCATAAG). After transfection and flow sorting of green fluorescent cells, clonal lines with the lowest inward current amplitude determined by whole-cell automated patch clamp recording were selected.

### Immunoblotting

Mouse and rat Na_V_1.6 and Na_V_1.7 were detected using anti-Na_V_1.6 antibody (1:100 dilution, ASC-009, Alomone Labs, Jerusalem, Israel) and anti-Na_V_1.7 antibody (1:200 dilution, 75-103, UC Davis/NIH NeuroMab Facility, Davis, CA), respectively. Transferrin (loading control) was detected with mouse anti-human transferrin receptor antibody (1:500, Cat# 136800, Invitrogen, Waltham, MA). Protein samples (50 µg) were electrophoresed on 7.5% polyacrylamide Mini-PROTEAN TGX gels (4561023, Bio-Rad, Hercules, CA), electrotransferred to Immobilon_®_-P PVDF Membrane (pore size 0.45um, IPFL00010, Millipore Sigma), and blocked in 5% bovine serum albumin for 1 hour. Sodium channels were detected by first incubating with Na_V_1.6 or Na_V_1.7 primary antibodies followed by incubation with IRDye 800CW goat anti-mouse antibody (1:10,000, catalog # 926-322100, LI-COR, Lincoln, NE). Transferrin was detected with primary antibody followed by incubation with IRDye 680RD goat anti-mouse antibody (1:10,000, catalog #926-68070, LI-COR, Lincoln, NE). Protein bands were imaged with the Li-Cor Odyssey CLx Imaging System (Li-Cor Biosciences, Lincoln, NE, USA).

### Transfections

Wild-type and variant human Na_V_1.6A and Na_V_1.6N plasmids were transiently expressed in ND7/LoNav cells using the Maxcyte STX system (MaxCyte Inc., Gaithersburg, MD, USA). ND7/LoNav cells were seeded at 1.1×10_6_ cells/ml in 100 mm tissue culture dishes and grown to 65-75% density then harvested using TrpLE_TM_ Express (GIBCO Cat. #12605010, Thermo Fisher Scientific, Waltham, MA, USA). A 500 µl aliquot of cell suspension was used to determine cell number and viability using an automated cell counter (ViCell, Beckman Coulter, Indianapolis, IN, USA). Remaining cells were collected by gentle centrifugation (160 × g, 4 minutes), washed with 5 ml electroporation buffer (EBR100, MaxCyte Inc.), and re-suspended in electroporation buffer at a density of 10^8^ viable cells/ml. Each electroporation was performed using 100 µl of cell suspension.

ND7/LoNav cells were electroporated with 50 µg of WT or variant Na_V_1.6 cDNA. The DNA-cell suspension mix was transferred to an OC-100 processing assembly (MaxCyte Inc.) and electroporated using the preset Optimization 4 protocol. Immediately after electroporation, 10 µl of DNase I (Sigma-Aldrich, St. Louis, MO, USA) was added to the DNA-cell suspension. Cell-DNA-DNase mixtures were transferred to 6-well tissue culture plate and incubated for 30 min at 37°C in 5% CO_2_. Following incubation, cells were gently re-suspended in culture media, transferred to a 100 mm tissue culture dish and grown for 48 hours at 37°C in 5% CO_2._ Following incubation, cells were harvested, counted, transfection efficiency determined by flow cytometry (see below), and then frozen in 1 ml aliquots at 1.8×10_6_ viable cells/ml in liquid nitrogen until used in experiments.

Transfection efficiency was evaluated by flow cytometry using a FacsCanto (BD Biosciences, Franklin Lake, NJ) located in the Northwestern University Interdepartmental Immunobiology Flow Cytometry Core Facility. Forward scatter, side scatter, and red fluorescence were measured using a 488 nm laser.

### Manual patch clamp recording

Currents were recorded at room temperature in the whole-cell configuration while acquired at 20 kHz and filtered at 5 kHz. Bath solution contained (in mM): 145 NaCl, 4 KCl, 1.8 CaCl_2_, 1 MgCl_2_, 10 HEPES ((*N*-(2-hydroxyethyl)piperazine-*N′*-2-ethanosulphonic acid), pH 7.35, and 310 mOsm/kg. The composition of the pipette solution was (in mM): 10 NaF, 110 CsF, 20 CsCl, 2 EGTA (ethylene glycol-bis-(β-aminoethyl ether)-tetraacetic acid), 10 HEPES, pH 7.35, 310 mOsm/kg. Whole-cell patch pipettes were pulled from thin-wall borosilicate glass (Warner Instruments, LLC, Hamden, CT, USA) with a multistage P-97 Flaming-Brown micropipette puller (Sutter Instruments Co., San Rafael, CA, USA) and fire-polished with a Micro Forge MF 830 (Narashige International, Amityville, NY, USA). Pipette resistance was ∼2 MΩ.

### Cell preparation for automated electrophysiology

Electroporated cells were thawed the day before experiments, plated in 60 mm tissue culture dishes and incubated for 18-24 hours at 37°C in 5% CO_2_. Prior to experiments, cells were detached using TrpLE™Express, resuspended in cell culture media and counted. Cells were centrifuged at 100 × g for 2 minutes then resuspended at 180,000/ml with external solution (see below) and allowed to recover 45 minutes at 15°C while shaking 200 rpm on a rotating platform.

### Automated Patch Clamp

Automated planar patch clamp recording was performed using a SyncroPatch 768 PE (Nanion Technologies) as previously described (23). External solution contained (in mM): 140 NaCl, 4 KCl, 2.0 CaCl_2_, 1 MgCl_2_, 10 HEPES, 5 glucose pH 7.4. The composition of the internal solution was (in mM): 10 NaF, 110 CsF, 10 CsCl, 20 EGTA, 10 HEPES, pH 7.2. Whole-cell currents were acquired at 10 kHz and filtered at 3 kHz. The access resistance and apparent membrane capacitance were determined. Series resistance was compensated 90% whereas leak and capacitance artifacts were subtracted using the P/4 method.

Data were analyzed and plotted using a combination of DataController384 version 1.8 (Nanion Technologies), Excel (Microsoft Office 2013, Microsoft), SigmaPlot 2000 (Systat Software, Inc., San Jose, CA USA) and Prism 8 (GraphPad Software, San Diego, CA) as previously described (23). Whole-cell currents were measured from a holding potential of -120 mV. Whole-cell conductance (G_Na_) was calculated as G_Na_ = *I*/(*V* − *E*_*rev*_), where *I* is the measured peak current, *V* is the step potential, and *E*_*rev*_ is the calculated sodium reversal potential. G_Na_ at each voltage step was normalized to the maximum conductance between −80 mV and 20 mV. To calculate voltage dependence of activation and inactivation, data were plotted against voltage and fitted with Boltzmann functions. Time-dependent recovery from inactivation was evaluated by fitting peak current recovery with a two-exponential function. Time-dependent entry into inactivation was evaluated by fitting current decay at 0 mV with a single exponential function. Number of cells (n) is given in the figure legends. Persistent current was measured as the ratio of peak ramp current to the peak current measured at 0 mV during the activation protocol; and charge movement was calculated as the integral of the ramp current divided by the current measured during the activation protocol. Window current was calculated by integrating the area under the intersection between the Boltzmann fits for voltage-dependence of activation and inactivation using a custom MatLab script (24).

### Statistics

Data from individual cells expressing variant Na_V_1.6 were compared to the average of the isoform-matched WT Na_V_1.6 run in parallel using a t-test. Statistical significance was established at P ≤ 0.01, which represents a Bonferroni’s correction for multiple testing in experiments comparing 4 variants to WT channels. One-way ANOVA was used to compare functional properties of WT Na_V_1.6 splice isoforms and non-transfected cells with statistical significance set at P ≤ 0.01. Unless otherwise noted, data are presented as mean ± 95% confidence intervals (CI).

## Data availability

The authors confirm that the data supporting the findings of this study are available within the article and its Supplementary material.

## AUTHORS’ CONTRIBUTIONS

C.G.V., T.V.A., J.M.D., N.F.G., M.J.O., C.B.B., and C.H.T. performed experimental work and analyzed data. I.H. acquired informed consent. A.L.G. acquired funding and supervised the work. The manuscript was written primarily by C.G.V. and A.L.G. with input from all co-authors. All authors reviewed and approved the final version of the manuscript prior to submission.

## ACKNOWLEDGEMENTS

This study was funded by a grant from the National Institute for Neurological Diseases and Stroke (NS108874).

## DISCLOSURES

A.L.G. received research grant funding from Praxis Precision Medicines, Neurocrine Biosciences, and Biohaven Pharmaceuticals, and received research grant funding from and is a member of the Scientific Advisory Board for Tevard Biosciences.

## SUPPLEMENTAL INFORMATION

### Supplemental Figures

**Fig. S1.**
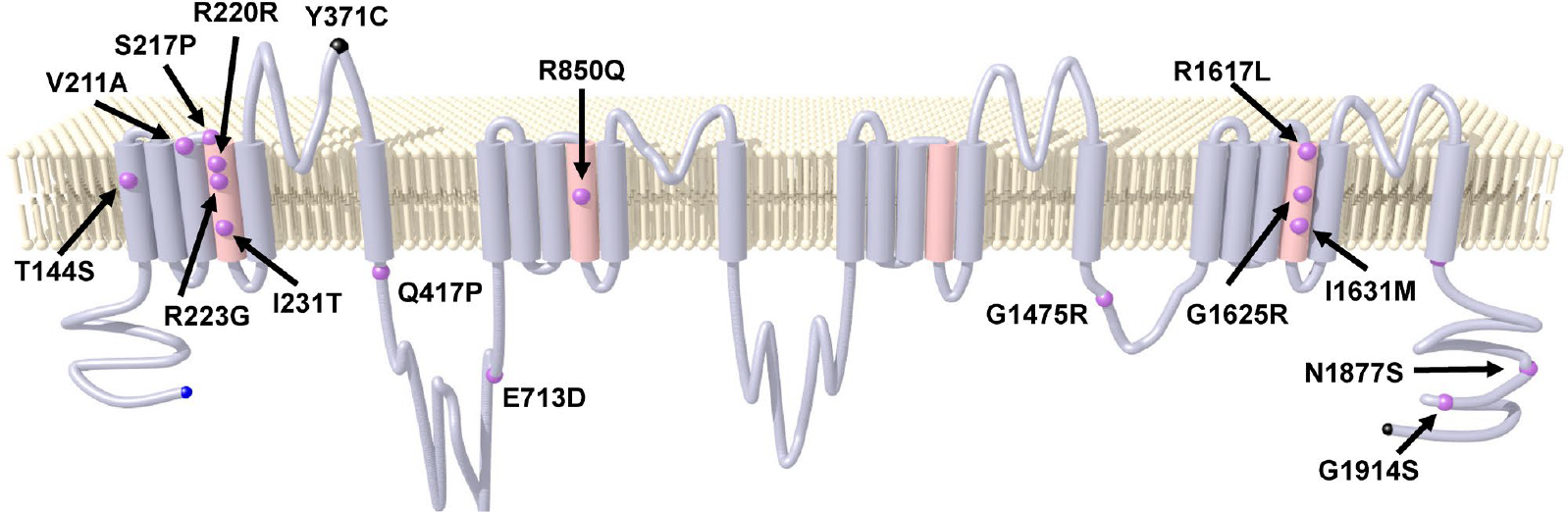
Location of Na_V_1.6 variants analyzed in this study. Simplified transmembrane topology of Na_V_1.6 with locations of variants studied.

**Fig. S2.**
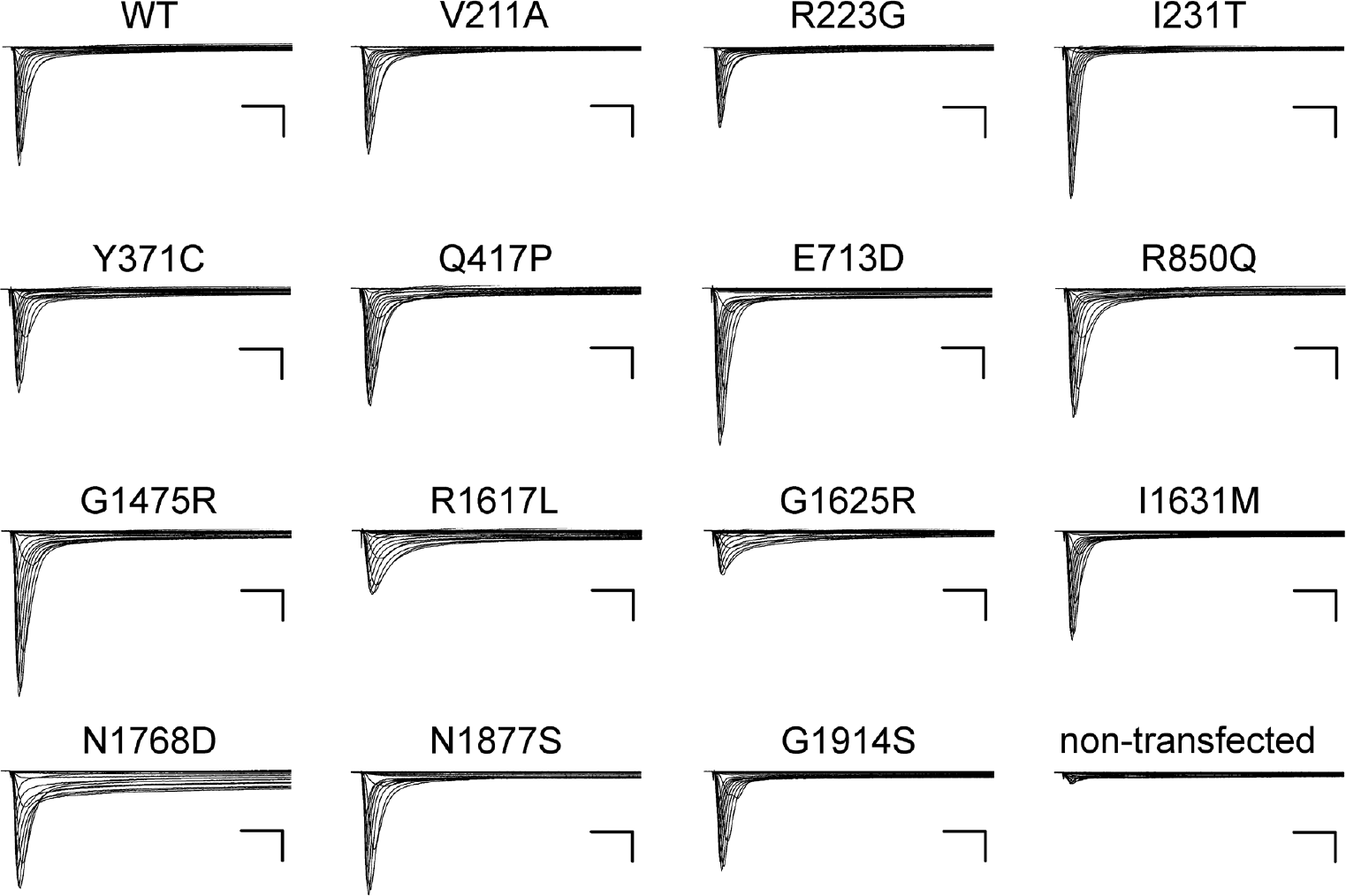
Whole-cell currents for variants expressed in Na_V_1.6A. Averaged whole-cell currents recorded from ND7/LoNav cells transfected with Na_V_1.6 variants and normalized to the WT channel peak current recorded in parallel (n = 27-64 per variant). Scale bars are 5 ms (horizontal) and 25% of WT channel current density (vertical).

**Fig. S3.**
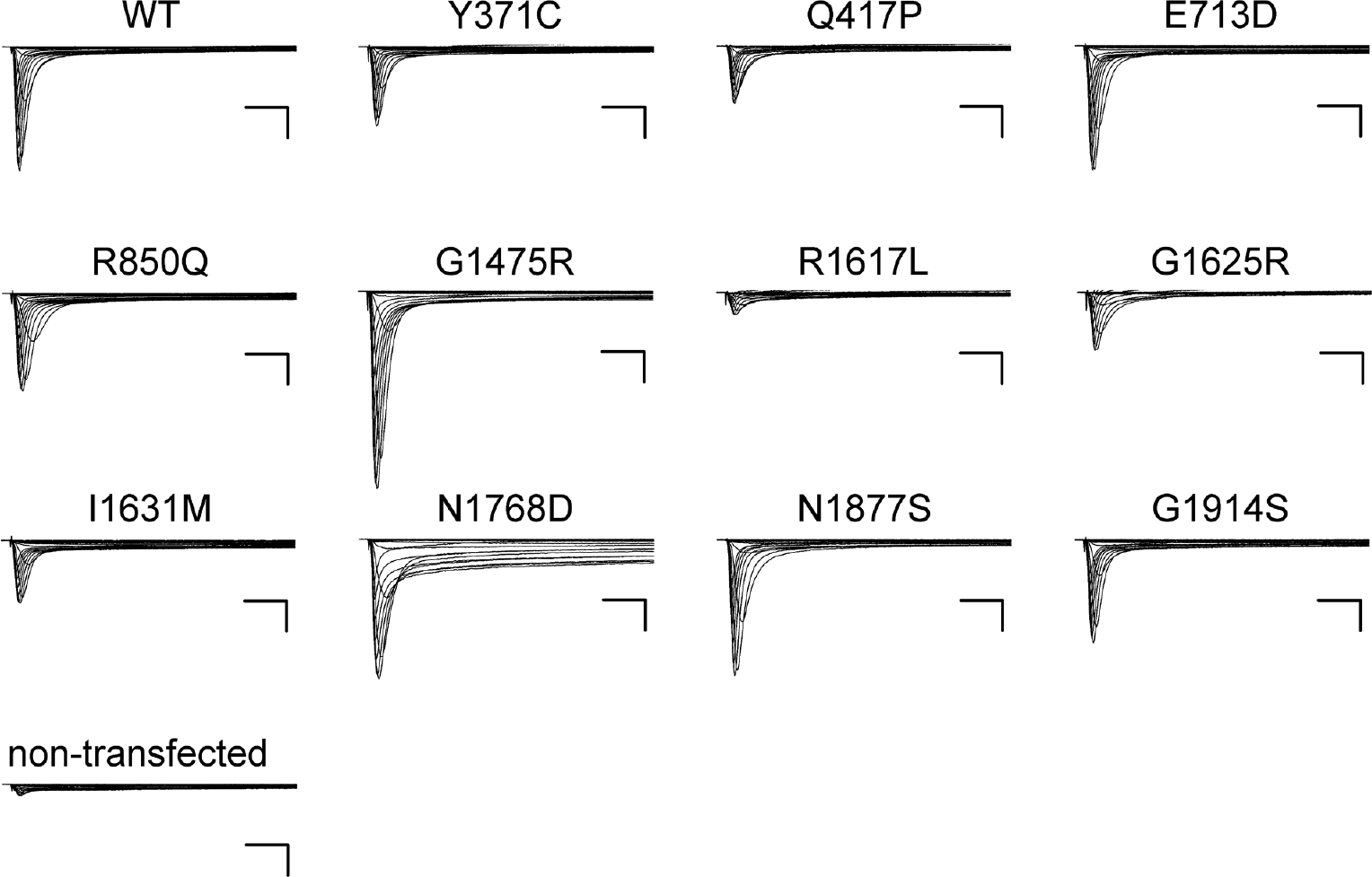
Whole-cell currents for variants expressed in Na_V_1.6N. Averaged whole-cell currents recorded from ND7/LoNav cells transfected with Na_V_1.6 variants and normalized to the WT channel peak current recorded in parallel (n = 25-67 per variant). Scale bars are 5 ms (horizontal) and 25% of WT channel current density (vertical).

**Fig. S4.**
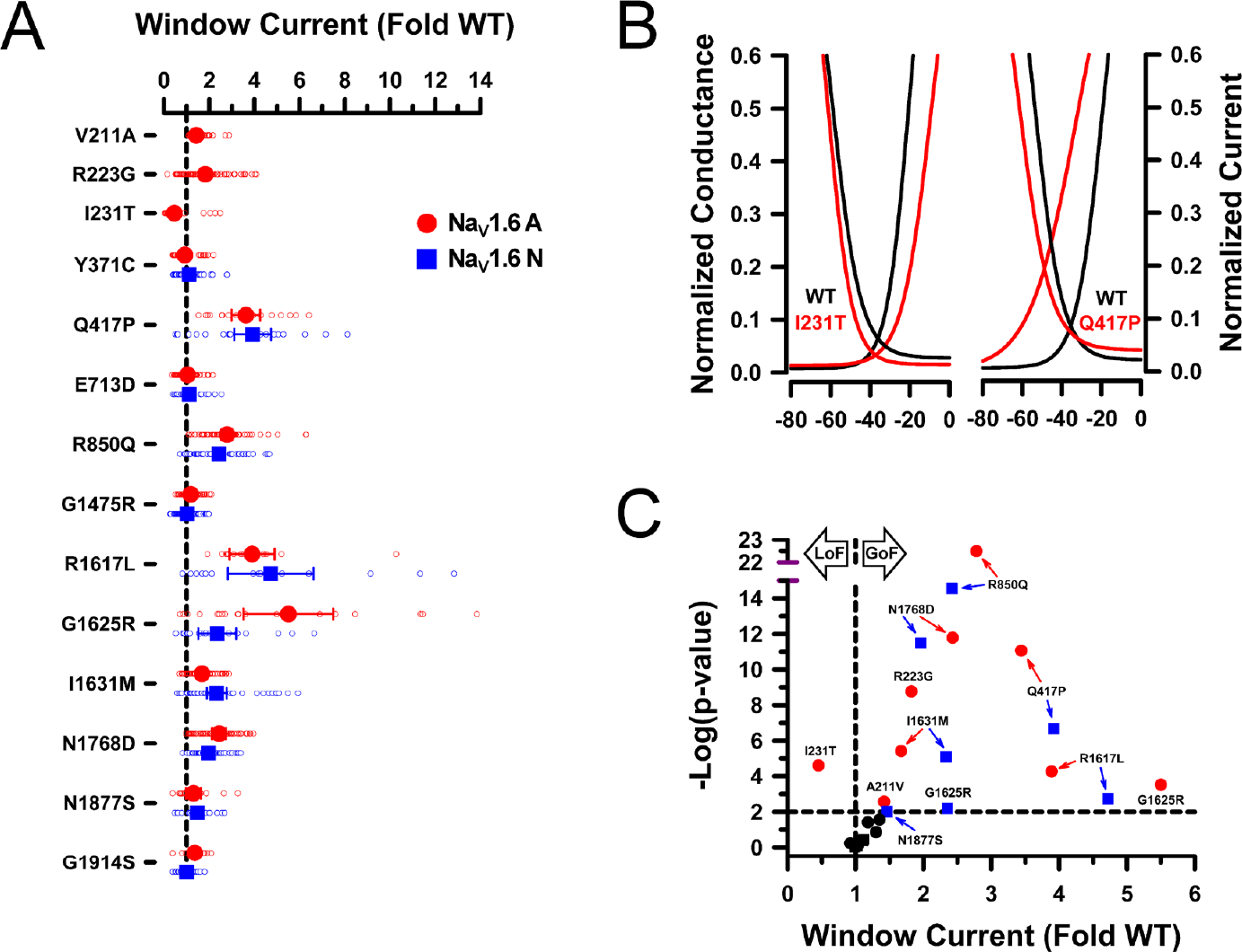
Window current determined for Na_V_1.6 variants. (**A**) Average deviation from WT Na_V_1.6 for window current area. All individual data points are plotted as open symbols and mean values are shown as larger filled symbols (n = 14-69). Error bars represent 95% CI. Data from Na_V_1.6A or Na_V_1.6N are indicated as red or blue symbols, respectively. Values to the right or left of the vertical dashed line (average normalized WT value) indicate larger (gain-of-function) or smaller (loss of function) window current, respectively. (**B**) Averaged Boltzmann fit lines for activation and steady-state inactivation curves of representative variants with smaller (I231T) or larger (Q417P) window current. (**C**) Volcano plot of mean values highlighting variants significantly different (P<0.01, horizontal dashed line) from WT. Symbols to the right of the vertical dashed line represent larger window current (gain-of-function). Only one variant (I231T) exhibited significantly smaller window current. Black symbols represent variants with no significant difference from WT. Quantitative data with statistical comparisons are provided in **Supplemental Dataset S2** (Na_V_1.6N) and **Supplemental Dataset S3** (Na_V_1.6A).

**Fig. S5.**
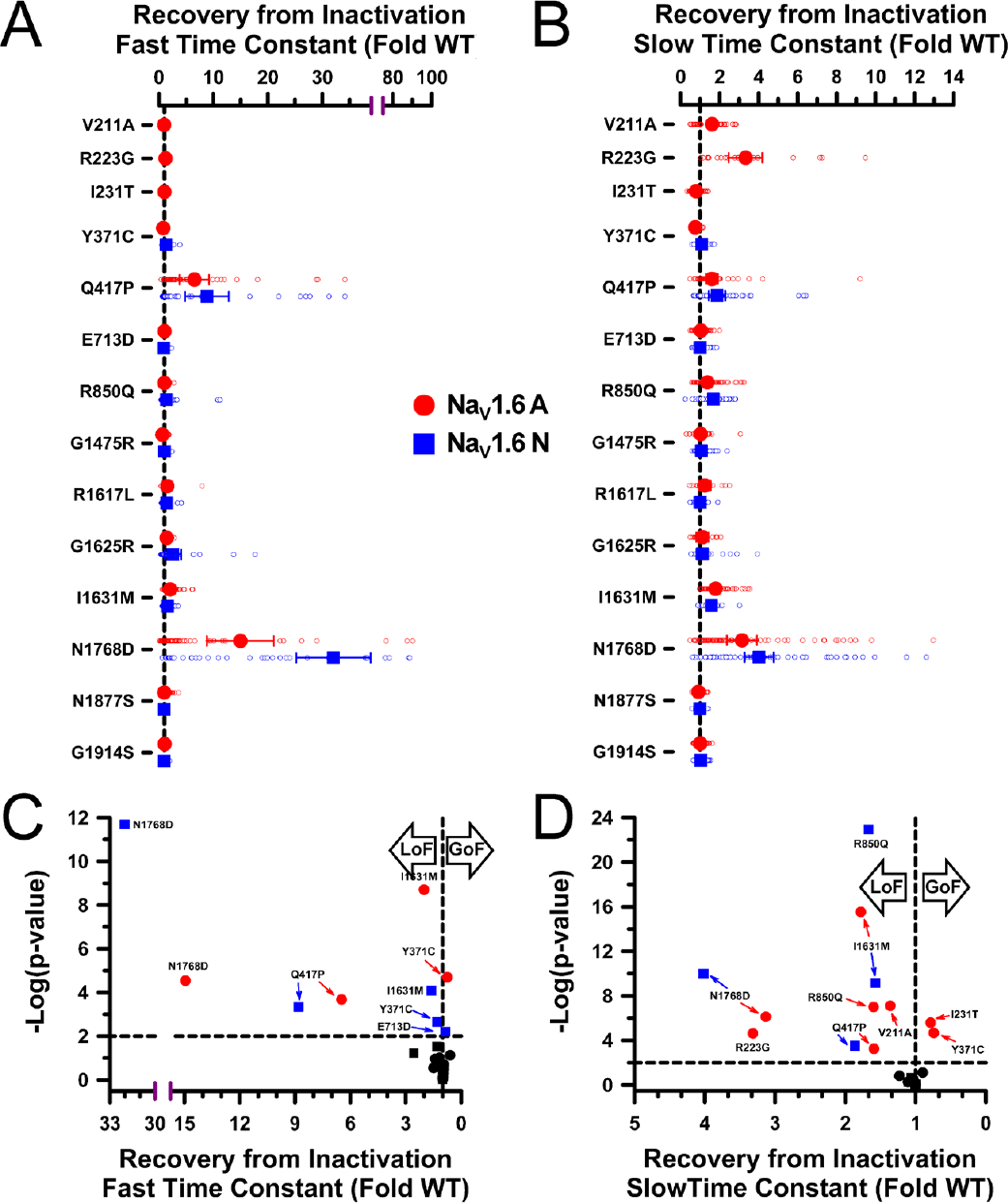
Recovery from inactivation determined for Na_V_1.6 variants. **(A**,**B)** Averaged time constants for recovery from inactivation displayed as fold-difference from WT channels recorded in parallel. Time constants (fast component plotted in panel A, slower component plotted in panel B) were determined by fitting the time course of recovery from inactivation to a double exponential function. All individual data points are plotted as open symbols and mean values are shown as larger filled symbols (n = 12-105 per variant). Error bars represent 95% confidence intervals. Data from Na_V_1.6A or Na_V_1.6N are indicated as red or blue symbols, respectively. Values to the right or left of the vertical dashed line (normalized WT value) represent larger (slower recovery) or smaller (faster recovery) time constants, respectively. **(C**,**D)** Volcano plots highlighting variants with significantly different (P <0.01, horizontal dotted line) fast component **(C)** and slower component **(D)** time constants of recovery from inactivation. Symbols to the left of the vertical dotted line denote slower recovery time course (loss-of-function), while symbols to right indicate faster recovery time course (gain-of-function). Black symbols represent variants with no significant difference from WT. Quantitative data with statistical comparisons are provided in **Supplemental Dataset S2** (Na_V_1.6N) and **Supplemental Dataset S3** (Na_V_1.6A).

**Fig. S6.**
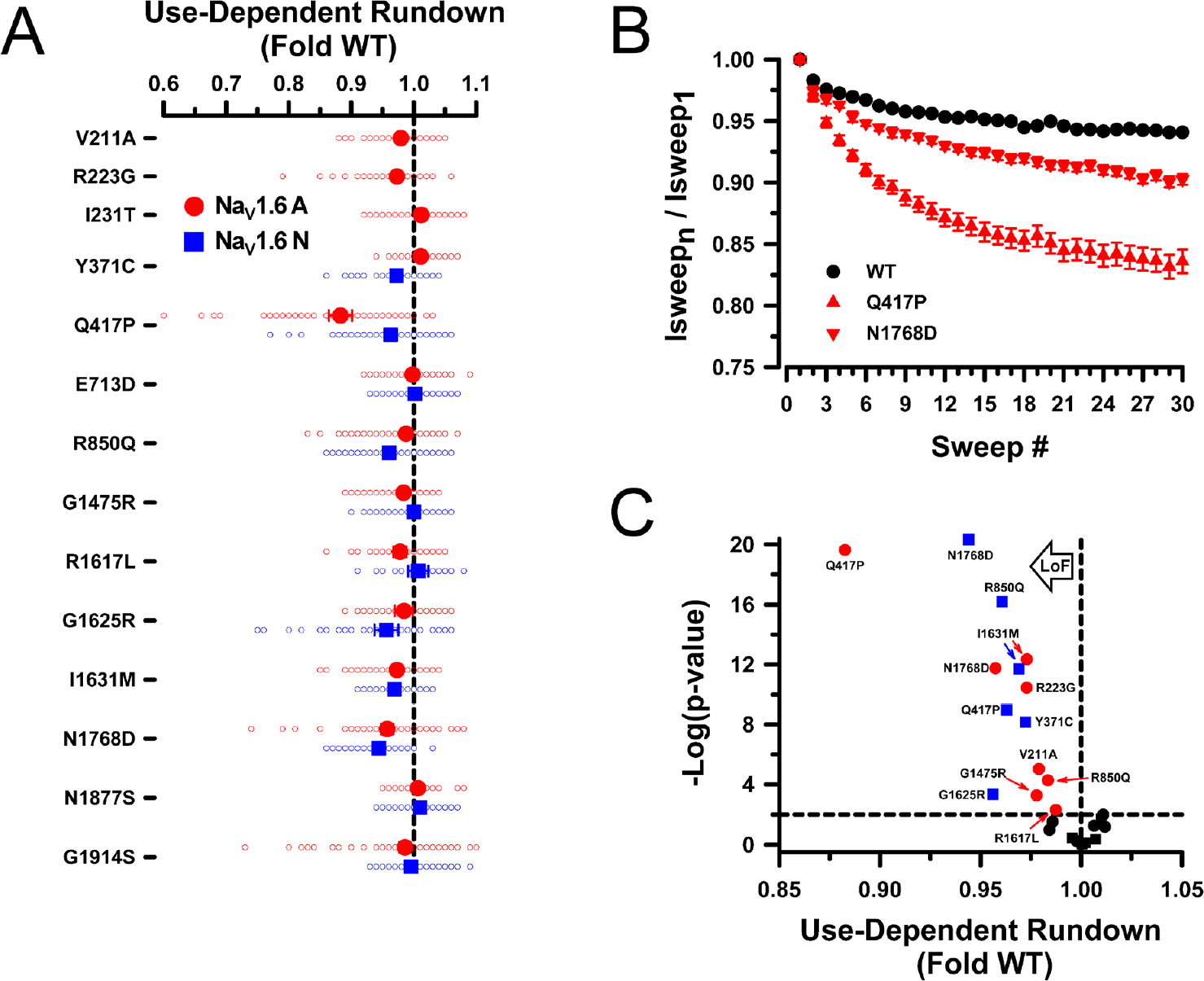
Use-dependent rundown of Na_V_1.6 variants. **(A)** Averaged use-dependent channel rundown at 20 Hz measured for Na_V_1.6 variants displayed as fold-difference from WT channels recorded in parallel. All individual data points are plotted as open symbols and mean values are shown as larger filled symbols (n = 26-153 per variant). Error bars represent 95% confidence intervals. Data from Na_V_1.6A or Na_V_1.6N are indicated as red or blue symbols, respectively. Values to the left of the vertical dashed line (normalized WT value) represent greater rundown than WT. No variants exhibited lesser degree of rundown than WT. **(B)** Averaged currents normalized to first sweep amplitude measured for 30 sweeps at 20 Hz for select variants expressed in Na_V_1.6A. **(C)** Volcano plot of mean values highlighting variants with significantly different (P<0.01, horizontal dashed line) use-dependent rundown from WT. Symbols to the left of the vertical dashed line represent greater rundown (loss-of-function and there were no variants with lesser rundown. Black symbols represent variants with no significant difference from WT. Quantitative data with statistical comparisons are provided in **Supplemental Dataset S2** (Na_V_1.6N) and **Supplemental Dataset S3** (Na_V_1.6A).

**Fig. S7.**
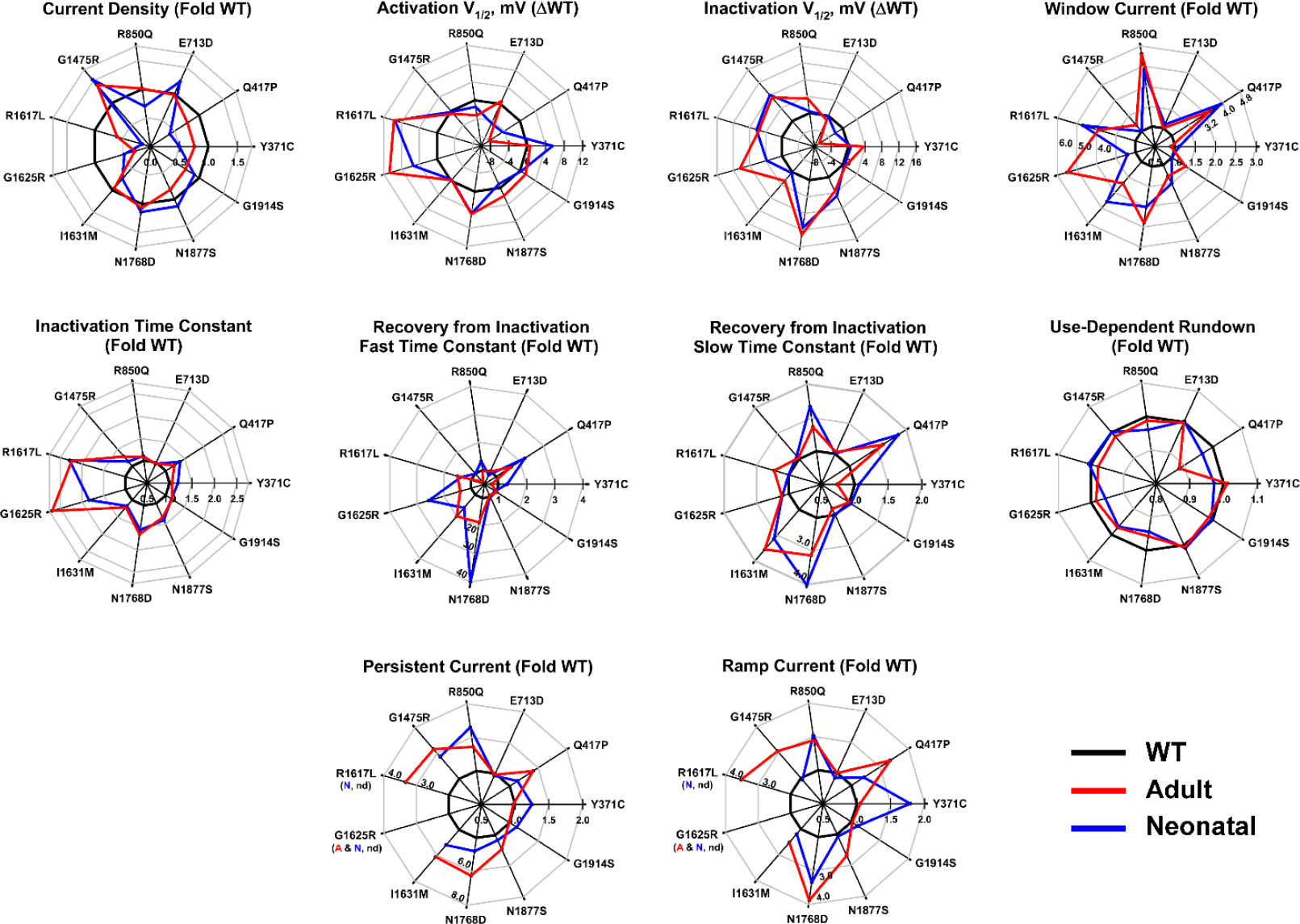
Comparison of functional properties among SCN8A variants. Radar plots depicting biophysical properties among variants compared to the WT channel. Individual radar plots represent different properties and each individual variant is represented as points (mean values normalized to WT) along each spoke. Red lines connecting each point represent data from Na_V_1.6A, blue lines represent data from Na_V_1.6N, and black lines indicate isoform-matched WT values. The scale indicating the magnitude of difference for each biophysical property is shown within the radar plot on the Y371C spoke, except for individual variants and specific properties (e.g., Window Current for Q417P and R1617L; Persistent Current for R1617L and N1768D; and Ramp Current for R1617L and N1768D). Persistent Current and Ramp Current were not determined (nd) for some variants. Quantitative data with statistical comparisons are provided in **Supplemental Dataset S2** (Na_V_1.6N) and **Supplemental Dataset S3** (Na_V_1.6A).

**Fig. S8.**
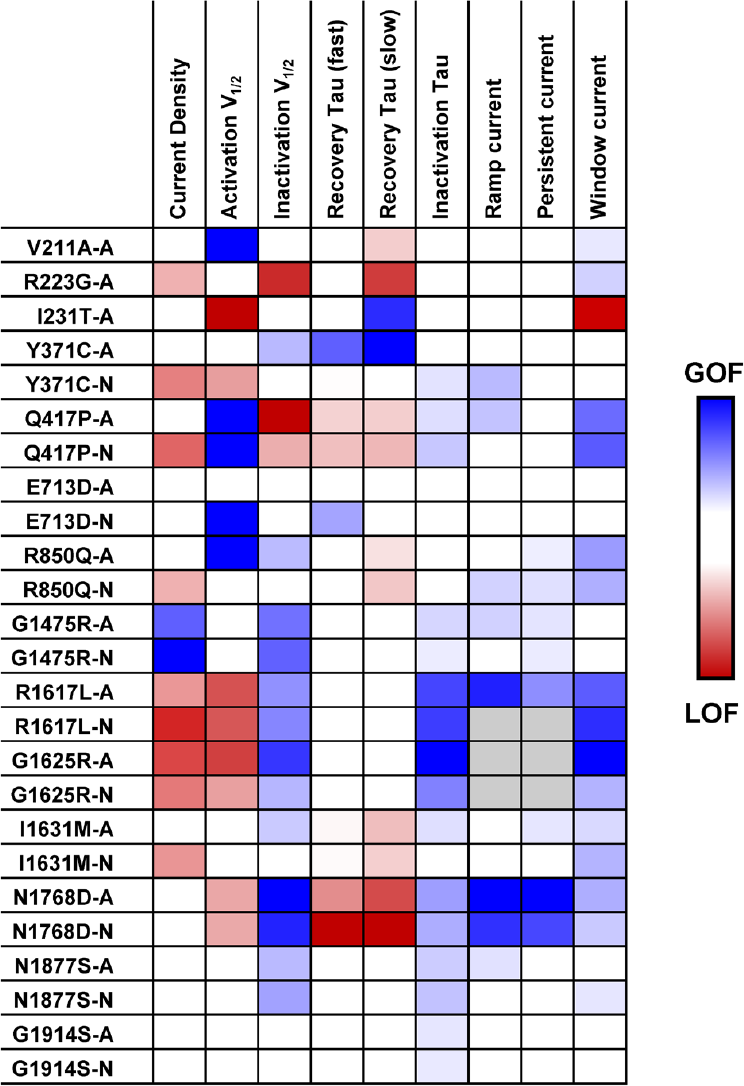
Heat map illustrating functional properties of Na_V_1.6 variants. Summary of individual functional properties for variants expressed in either Na_V_1.6A or Na_V_1.6N are illustrated as a heat map. Properties with gain-of-function are shaded as blue, and loss-of-function as red. Gray shaded boxes are properties that were not determined, and uncolored boxes indicate WT-like values. Only properties that reached the threshold for statistical significance (P<0.01) are color coded. The intensity of shading reflects the degree of difference with WT.

### Supplemental Tables

**Table S1.** Phenotypes of *SCN8A* variants in this study.

| Nucleotide | Protein | Recurrent | Phenotype | Age at onset | Reference(s) |
| --- | --- | --- | --- | --- | --- |
| c.632T>C | V211A | N | Epileptic encephalopathy | 3 m | Exon 5A: (1)<br>Exon 5N: (2) |
| c.667A>G | R223G | N | West syndrome | 4-6 m | Exon 5A: (1)<br>Exon 5N: (3)<br>Unclear: (4) |
| c.692T>C | I231T | N | Epileptic encephalopathy | 7-8 m | Exon 5N: (1) |
| c.1250A>C | Q417P | N | Intractable infantile spasms, global developmental delay | < 12 m | This study and (2) |
| c.2139A>C | E713D | N | Variant of uncertain significance | Unknown | ClinVar ID 838164 |
| c.2549G>A | R850Q | Y | Epileptic encephalopathy | 2 m | (5-8) |
| c.4423G>A | G1475R | Y | Epileptic encephalopathy, intermediate severity | 1 – 11 m | (6-13) |
| c.4850G>T | R1617L | Y | Early-onset epilepsy | 3-6 m | (5, 7, 8, 10, 12, 14, 15) |
| c.4893C>G | I1631M | N | Post-vaccination generalized seizures, later-onset partial epilepsy, mild intellectual disability | 5 m | This study |
| c.4873G>A | G1625R | Y | Intellectual disability, developmental delay | unknown | (16) |
| c.5302A>G | N1768D | Y | Epileptic encephalopathy, SUDEP | 6 m | (17) |
| c.5630A>G | N1877S | Y | Self-limited infantile epilepsy | 5 m | (18, 19) |
| c.5740G>A | G1914S | N | Variant of uncertain significance | unknown | ClinVar ID 2500085 |

**Table S2.**
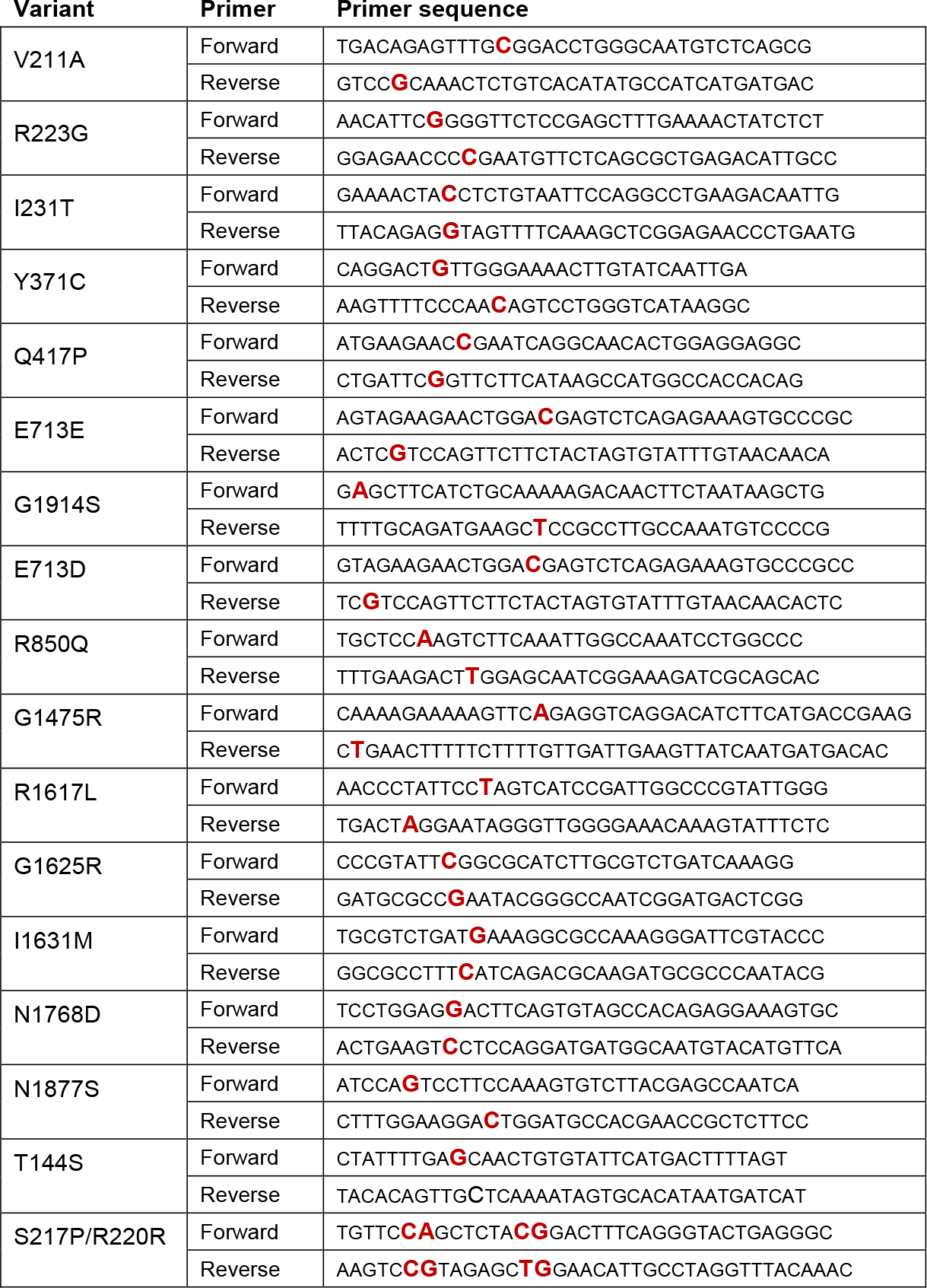
Mutagenic primer sequences for *SCN8A* variants (mutations are in bold red letters)

### Supplemental References

### Supplemental Datasets (contact authors)

Dataset S1. Biophysical properties of WT Na_V_1.6A and WT Na_V_1.6N (Excel file).

Dataset S2. Biophysical properties of variants studied in Na_V_1.6N (Excel file).

Dataset S3. Biophysical properties of variants studied in Na_V_1.6A (Excel file).

